# Signatures of plant defense response specificity mediated by herbivore-associated molecular patterns in legumes

**DOI:** 10.1101/2021.09.02.458788

**Authors:** Adam D. Steinbrenner, Evan Saldivar, Nile Hodges, Antonio F. Chaparro, Eric A. Schmelz

## Abstract

I.

Chewing herbivores activate plant defense responses through a combination of mechanical wounding and elicitation by herbivore associated molecular patterns (HAMPs). HAMPs are wound response amplifiers; however, specific defense outputs may also exist that strictly require HAMP-mediated defense signaling. To investigate HAMP-mediated signaling and defense responses, we characterized cowpea transcriptome changes following elicitation by inceptin, a peptide HAMP common in Lepidoptera larvae oral secretions. Following inceptin treatment, we observed large-scale reprogramming of the transcriptome consistent with 3 different response categories: 1) amplification of mechanical wound responses, 2) temporal extension through accelerated or prolonged responses, and 3) examples of inceptin-specific elicitation and suppression. At both early and late timepoints, namely 1 and 6 hours, large sets of transcripts specifically accumulated following inceptin elicitation but not wounding alone. Further inceptin-regulated transcripts were classified as reversing changes induced by wounding alone. Within key signaling and defense related gene families, inceptin-elicited responses commonly targeted select subsets of wound-induced transcripts. Transcripts displaying the largest inceptin-elicited fold-changes included terpene synthases (TPS) and peroxidases (POX) that correspond with induced volatile production and increased peroxidase activity in cowpea. Characterization of inceptin-elicited cowpea defenses via heterologous expression in *Nicotiana benthamiana* demonstrated that specific cowpea TPS and POX were able to confer terpene emission and the reduced growth of beet armyworm (*Spodoptera exigua*) herbivores, respectively. Collectively, our present findings in cowpea support a model where HAMP-elicitation both amplifies concurrent wound responses and specifically contributes to the activation of selective outputs associated with direct and indirect anti-herbivore defenses.

**Significance Statement:** Plants recognize herbivore-associated molecular patterns (HAMPs) to induce defenses, but interactions with the more general wound response are not well-understood. We leveraged a known HAMP-receptor interaction to characterize transcriptomic modulation of the wound response by the peptide HAMP, inceptin. Inceptin not only amplifies wound responses, but can specifically induce or suppress transcripts with demonstrated functions in direct and indirect defense against herbivores. The plant immune system thus recognizes HAMPs to fine-tune wound responses against herbivory.

## III. Introduction

Plants undergo large-scale physiological changes following attack by insect herbivores (Karban and Baldwin, 2007). Herbivore-induced plant responses include direct defenses such as enhanced production of antinutritive proteins, toxic or deterrent non-volatile specialized metabolites and indirect defenses such as low molecular weight volatiles that attract natural enemy bodyguards from higher trophic levels (Howe and Jander, 2008). Insect herbivory includes different forms of physical wounding, ranging from targeted cellular damage by piercing/sucking insects to large-scale tissue removal by mandibles and chewing. Wound- and herbivore-induced plant responses are mediated by a large set of defense-related phytohormones, most commonly examined as the interplay of jasmonates, ethylene, and salicylic acid (Schmelz *et al*., 2007) that mediate tradeoffs with growth-related processes (Pieterse *et al*., 2012).

Despite shared signaling components, plant responses to either wounding or live herbivory are qualitatively different, and in many cases wounding alone is unable to recapitulate the full set of induced plant responses to herbivory (Bricchi *et al*., 2010; Li *et al*., 2019; Duran-Flores and Heil, 2016; Schmelz *et al*., 2009; Schmelz *et al*., 2006; Musser *et al*., 2002). Diverse factors associated with salivary secretions, glandular secretions, and regurgitant, collectively termed oral secretions (OS), modify plant responses to herbivory (Schmelz, 2015). OS contains both elicitors of defense, termed herbivore-associated molecular patterns (HAMPs), and suppressors, termed effector proteins (Chen *et al*., 2019; Musser *et al*., 2002). HAMPs are generally thought to amplify wound-induced signaling through specific recognition by the plant immune system (Stahl *et al*., 2018; Uemura and Arimura, 2019). HAMPs are conceptually analogous to well-studied pathogen- and damage-associated molecular patterns (PAMPs and DAMPs), which are critical for effective immune responses to bacterial, fungal, and oomycete pathogens (Gust *et al*., 2017; Couto and Zipfel, 2016). The adaptive balance of HAMP recognition by plants and suppression or evasion by herbivores likely contributes to highly specialized plant-herbivore interactions, similar to arms races of induced secondary metabolite production and counter-adaptations (Stahl *et al*., 2018).

Plant transcriptomic analyses following live insect herbivory have captured the large, combined effects of both wounding and OS (Heidel-Fischer *et al*., 2014; Reymond *et al*., 2000; Major and Constabel, 2006; De Vos *et al*., 2005; Qi *et al*., 2016; Appel *et al*., 2014), including potential suppressors in addition to HAMPs (Consales *et al*., 2012; Musser *et al*., 2002). In contrast, RNA-seq analyses focusing on the distinct contributions of both wounding and defined HAMPs remain uncommon. For example, a broadly occurring and widely active HAMP, namely the fatty acid-amino acid conjugate (FAC) N-linolenoyl-L-glutamine (18:3-Gln) elicits widespread transcriptional changes in *Nicotiana* sp. compared to wounded plants (Gilardoni *et al*., 2011; Xu *et al*., 2015), however, it is unclear which plant responses also occur following wounding alone.

In this study we utilize a biochemically defined HAMP, namely the chloroplastic ATP synthase γ-subunit protein (cATPC) fragment termed inceptin (Schmelz *et al*., 2006; Steinbrenner *et al*., 2020) to precisely define separate wound- and HAMP-elicited responses in a model legume. Among diverse HAMP signatures in Lepidopteran larval OS, inceptin-like peptides occur as a series of proteolytic fragments ranging from 10 to 13 amino acids long (Schmelz *et al*., 2007) and chemically vary based on the host plant cATP sequence (Schmelz *et al*., 2006). The most abundant inceptin variant in OS of caterpillar larvae feeding on cowpea (*Vigna unguiculata*) is the 11-amino acid inceptin (In) derivative ^(+^ICDINGVCVDA^-^), abbreviated *Vu-*In hereafter. Like many HAMPs and PAMPs (Schmelz *et al*., 2009; Boutrot and Zipfel, 2017), inceptin is only bioactive on a subset of plant species due to host variation in immune recognition. Related legume species in the Phaseolinae, but not any other tested plants, are able to respond to inceptin (Schmelz *et al*., 2009).

*Vu*-In interacts with wounding to activate several plant defense outputs. In previous studies, scratch-wounding cowpea leaves led to no significant induction of ethylene or (E)-4,8-dimethyl-1,3,7-nonatriene (DMNT) homoterpene volatiles, while addition of inceptin led to strong induced responses (Schmelz *et al*., 2006). Additional experiments using diet interventions demonstrated that the specific presence of inceptin in the OS is required for cowpea responses to short feeding bouts by live caterpillar herbivores (Schmelz *et al*., 2007). Wounding alone therefore is insufficient for full plant responses to herbivory in cowpea, and *Vu*-In serves as a critical HAMP elicitor to activate strong anti-herbivore defense responses.

At a molecular scale, plant recognition of both damage and specific HAMPs is mediated by an immune system of cell surface receptors detecting modified and non-self patterns, akin to pathogenic epitope PAMPs, and DAMPs such as self-derived cell wall fragments and peptide hormones (Gust *et al*., 2017; Boutrot and Zipfel, 2017). Unlike for PAMPs, until recently analogous plant perception mechanisms for HAMPs were unknown. We recently identified a specific cell surface receptor, termed the Inceptin Receptor (INR) that confers binding, signaling, and defense outputs in response to the peptide ligand inceptin (Steinbrenner *et al*., 2020). INR is a legume-specific leucine-rich repeat receptor-like protein. Expression of INR in non-native species such as tobacco allows *Vu*-In induced association with co-receptor kinases to activate canonical immune signaling and reduced armyworm caterpillar growth rates. Thus, empirical data supports the hypothesis that inceptin acts as a specific input signal to activate an INR-mediated response. We therefore predicted that immune signaling downstream of INR interacts with wounding to generate complex and unique *Vu*-In-specific defense outputs.

In this study, we characterized the transcriptome wide responses of cowpea leaves to both wounding and *Vu*-In treatment. First, we aimed to examine whether or not *Vu*-In interacts with wounding to activate specific responses or simply acts as an amplifier of wound responses. Second, we identified specific defensive processes modulated by the INR signaling pathway, including effects on signaling pathways, defense-related transcription factors (TFs), and metabolic outputs. Third, we sought to functionally validate proteins encoded by transcripts that dramatically accumulate following *Vu*-In elicitation to explore roles in direct and indirect defensive functions. Collectively, we show that the model HAMPs, such as inceptin, interact with the wound response to specifically modify expression of select gene families with anti-herbivore defense functions.

## IV. Results

### Wide-scale transcriptional wound responses in cowpea

To examine the interactive contribution of wounding and HAMP elicitation, leaves of intact cowpea plants were either 1) untreated or scratch-wounded with the additional application of either 2) H_2_O or 3) *Vu*-In peptide. After either 1 or 6 hr, scratch wounded leaves and adjacent untreated leaves were harvested. Unwounded leaf tissue harvested at 1 hr was required as a baseline for analysis of differentially expressed genes (DEG, see Materials and Methods for details). 10,365 genes showed significant differences between any two treatment groups (Table S1). Principal components analysis confirmed consistency of biological replicates across treatments and showed a strong effect of wounding and wounding plus inceptin treatment (Fig. 1A).

**Fig. 1.**
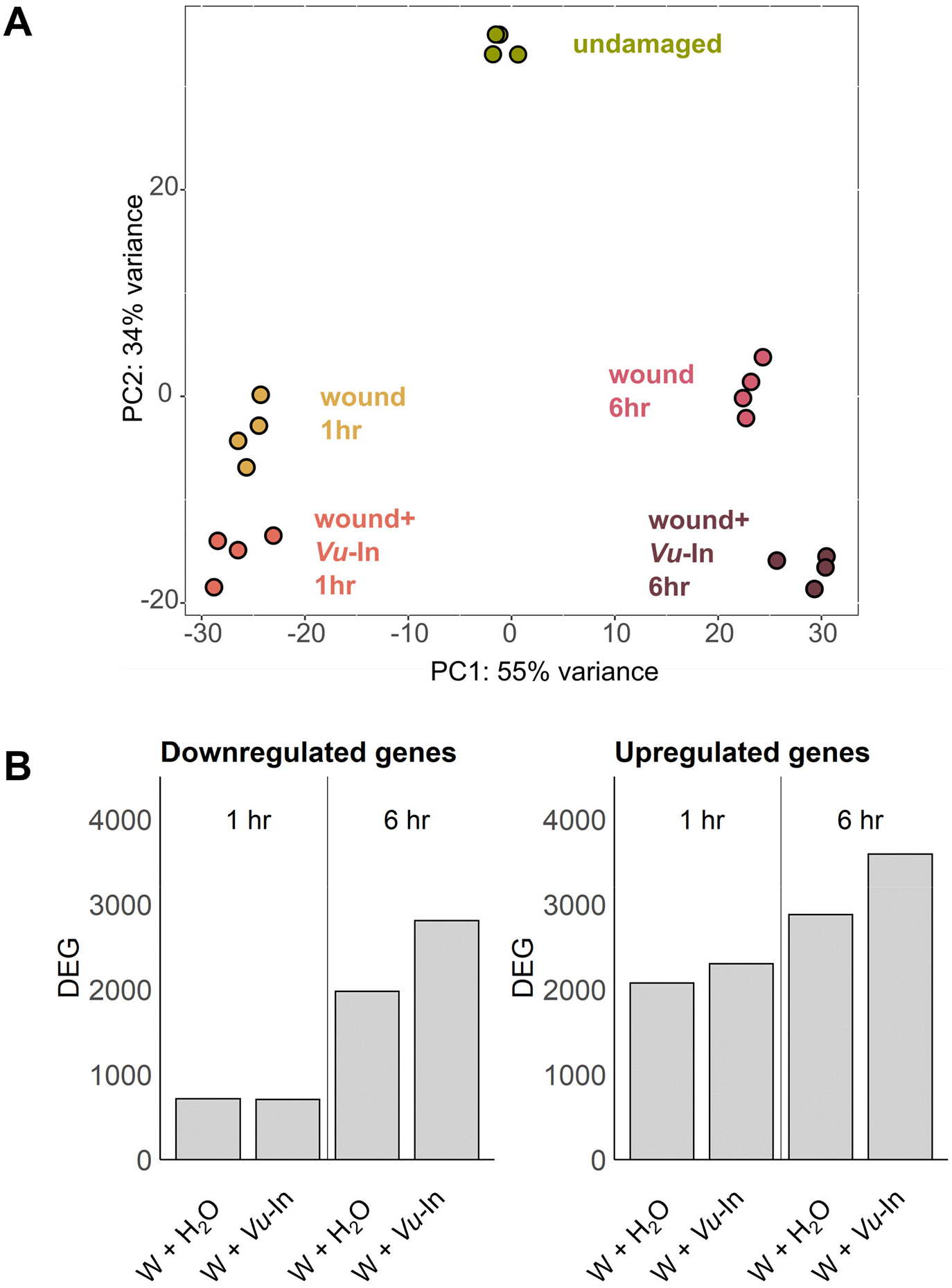
Transcriptomic analyses of wound and *Vu*-In elicited responses in cowpea. **A**, Principal component (PC) analysis of log-transformed counts data supports multiple treatment effects. Top two PC shown explain 89% of sample variance. **B**, Total number of differentially expressed genes (DEG) at 1 hr and 6 hr timepoints relative to undamaged tissue after wounding (W) or W + *Vu-*In treatment. Upregulated and downregulated DEGs are shown separately. DEG was defined as having abs(log2(FC)) > 2 relative to undamaged tissue and Benjamini-Hochberg adjusted p<0.05.

We next compared the sets of DEGs relative to unwounded tissue at both 1 and 6 hr (Fig. S1). In general, a larger set of DEGs was observed at the 6 hr timepoint (Fig. 1B), which may reflect genes under circadian control relative to the unwounded tissue harvested at 1 hr, as well as the effects of wounding. On top of these effects, *Vu-*In treatment increased the number of upregulated DEGs at both timepoints. In contrast, *Vu-*In treatment did not lead to a larger set of downregulated DEGs 1 hr after treatment (Fig. 1B). 73% and 83% of wound-induced gene expression changes also occurred after wound+inceptin treatment at 1 hr and 6 hr respectively (Fig. S2).

### Distinct transcript sets display HAMP specific regulation compared to wounding alone

We next analyzed differential expression between wound+*Vu*-In treatment compared to wounding alone. Spanning both the 1 and 6 hr time points, a total of 1443 transcripts were either significantly up- or down-regulated by *Vu*-In (Fig. 2A). To examine candidate induced defense responses, we separately analyzed up- and down-regulated transcripts. At 1 hr, 419 transcripts were significantly *Vu*-In-upregulated relative to wounding alone, and 109 were *Vu*-In -downregulated (Table S2). At 6 hr, 542 genes were inceptin-upregulated and 584 down-regulated (Table S3). The majority of both up- and down-regulated genes were unique to the individual timepoints (Fig. 2B), consistent with distinct early and late *Vu*-In-induced responses.

**Fig. 2.**
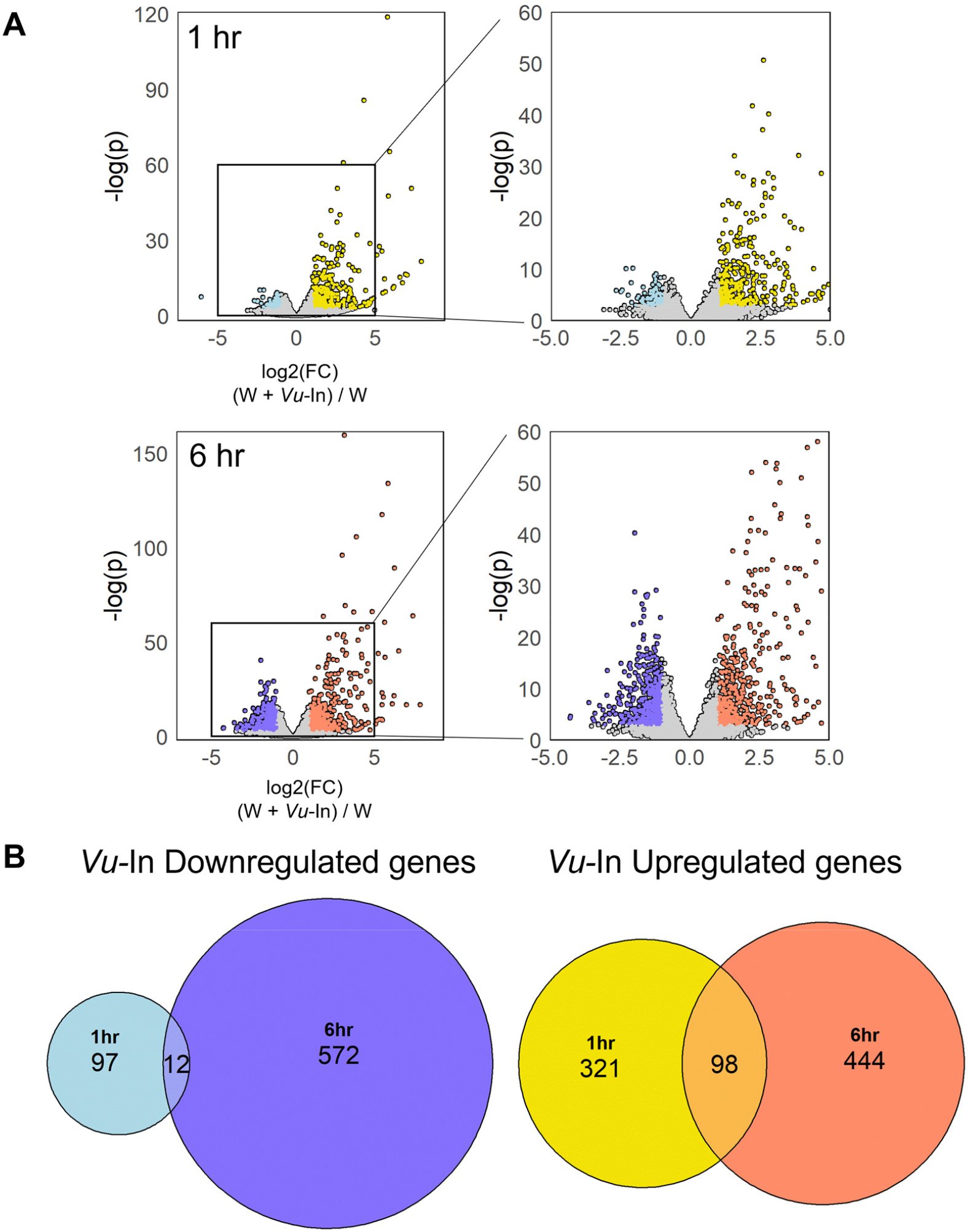
Early and late *Vu*-In-elicited transcriptional changes relative to parallel wound (W) controls at 1 hr and 6 hr. **A**, Volcano plot of *Vu-*In-elicited gene expression changes, plotting – log of adjusted p-value vs log2-corrected fold-change (FC). Left panels include all genes with detected expression, right panels are zoomed to identical axis scales across 1 hr and 6 hr. Significantly *Vu-*In elicited or –repressed genes are colored (abs(log2(FC)) > 1 and Benjamini-Hochberg adjusted p<0.05.) **B**, Venn diagrams of number of 1 hr vs 6 hr *Vu-*In elicited gene expression changes, with down- and upregulated genes shown separately. Cowpea gene identities underlying illustrated patterns are detailed in Tables S2 & S3.

To further understand these changes, we divided inceptin-induced genes into mutually-exclusive categories based on their relationship to wound-induced changes (Fig. S3). We filtered for three sets of genes: 1) *Vu*-In-amplified, 2) *Vu*-In-specific, and 3) *Vu*-In-accelerated/prolonged genes. At 1 hr, 193 genes were *Vu*-In-amplified, *i*.*e*. these genes were significantly upregulated relative to wounding alone, on top of significant upregulation by wounding relative to unwounded tissue (Fig. 3A, Table S2). The *Vu-*In-amplified gene set represented only 46% of total inceptin-induced genes at 1 hr, consistent with a complex interaction with the wound response.

**Fig. 3.**
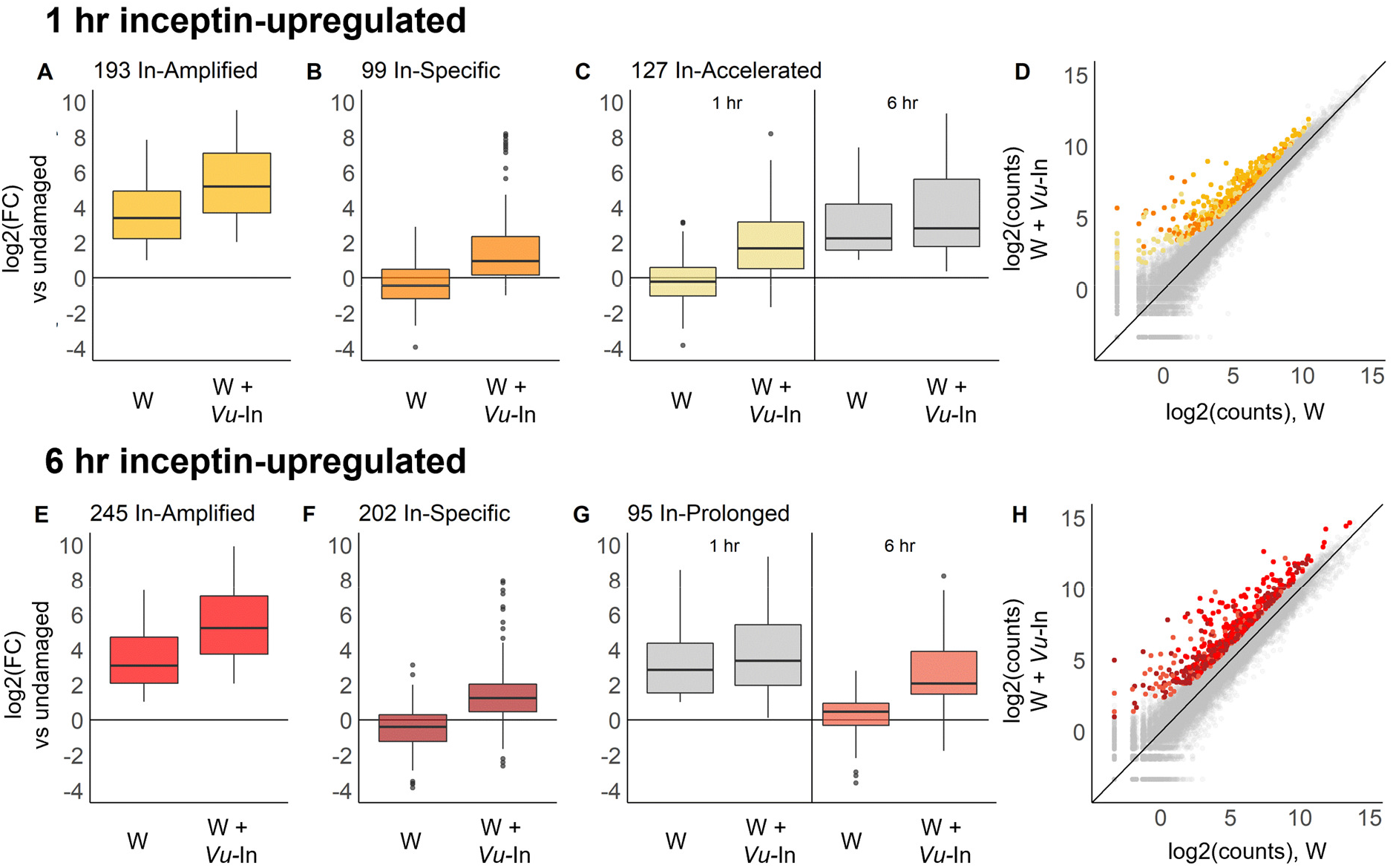
Transcriptional regulation of *Vu-*In-induced genes at 1 hr and 6 hr relative to wounding alone. Boxplots indicate magnitude fold change (FC) relative to undamaged treatment for categories of genes in **A**, *Vu*-In-Amplified; **B**, *Vu*-In-Specific; and **C**, *Vu*-In-Accelerated sets at 1 hr. **D**, Plot of log2(counts) after wounding (W) or W + *Vu-*In for genes in **A**-**C**. FC magnitude of **E**, *Vu*-In-Amplified; **F**, *Vu*-In-Specific; and **C**, *Vu*-In-Prolonged gene sets at 6 hr. **H**, Plot of log2(counts) after wounding (W) or W + *Vu-*In for differentially expressed genes in **E**-**G**. Summary of filtering criteria for the *Vu-In* induced category assignments for specific, amplified, accelerated and prolonged groups are detailed in Fig. S3. For panels C and G, the behavior of same set of genes is shown at both 1 hr and 6 hr. Cowpea genes in each category underlying illustrated patterns are detailed in Tables S2 & S3.

An additional 99 inceptin-specific genes were defined by significant upregulation at 1 hr but not by wounding alone (Fig. 3B). 53 of these 99 genes were in fact significantly reversed by inceptin relative to wound-downregulation (Table S4). Finally, 127 inceptin-accelerated genes were defined by their later induction at 6 hr by wounding alone (Fig. 1C, Table S2). Genes with diverse raw expression levels fell into all three categories (Fig. 3D). Rather than acting through simple wound amplification, *Vu*-In elicitation modulates the cowpea defense response.

Early 1 hr responses to *Vu-*In also included 109 down-regulated transcripts. Patterns varied markedly from the set of up-regulated transcripts. For example, in contrast to 193 *Vu-*In-amplified upregulated genes, no examples were detected with *Vu*-In further downregulating levels of a wound-downregulated transcript (Fig. S4A, Table S2). In contrast, 31 genes were inceptin-reversed, i.e. significantly downregulated by wounding but positively induced by inceptin treatment (Table S4). Inceptin therefore interacts with early wound-induced transcript repression exclusively as a positive regulator. 78 additional inceptin-induced changes represented either inceptin-specific changes (never observed in wound-only treatments) or inceptin-accelerated wound responses (Fig. S4B, S4C, Table S2).

At the later 6 hr timepoint, a largely distinct set of genes were induced by *Vu-*In (Fig. 2B). 542 genes were significantly upregulated relative to wounding alone, 245 of which were amplified wound responses (Fig. 3E, Table S3). In contrast, 202 changes were *Vu-*In-specific (Fig. 3F), and 95 showed a pattern of prolonged upregulation from damage-induced upregulation at the earlier timepoint (Fig. 3G). Genes with larger raw expression values were induced by inceptin at 6 hr vs 1 hr (Fig. 3H), reflecting large-scale transcriptional reprogramming. Similar to early responses, only a portion of inceptin-induced changes reflected amplification of wounding. Specifically 45% of the 6 hr *Vu*-In-induced genes fell into the *Vu*-In-amplified category.

Similar to late upregulated genes, only a portion of *Vu-*In-downregulated genes at 6 hr were categorized as amplification of wound-induced changes (152 of 584 of total downregulated DEGs, 26.0%) (Fig. S3, Table S3). Finally, inceptin treatment reversed wound-upregulated responses for 119 genes (Table S5).

### Inceptin modulates the accumulation of a subset of wound-induced transcripts encoding defense-related pathways

We next analyzed families of genes with functions in anti-herbivore signaling and defense. Generally, *Vu-*In-induced changes were greatly enriched for gene sets associated with defense related Gene Ontology (GO) terms, *e*.*g*. upregulation of isoprenoid and lipid biosynthesis and proteolysis, and downregulation of photosynthesis (Table S6). To understand these defense processes in more detail, we analyzed families of key defense-related genes in phylogenetic context with characterized Arabidopsis homologs. Consistent with the established *Vu*-In-mediated elicitation of ethylene and jasmonic acid production in cowpea (Schmelz *et al*., 2006; Schmelz *et al*., 2007), inceptin positively regulated gene sets encoding enzymes in both biosynthetic pathways (Fig. S5). Among ethylene biosynthetic genes, inceptin amplified the wound-induced expression of cowpea orthologs of Arabidopsis ACC synthase 6 and ACC oxidase 4 (Fig. S5A-C). Upregulation of jasmonate-related biosynthetic machinery, including 4 lipoxygenases and an allene oxide synthase, was either *Vu*-In-specific, -prolonged or -amplified compared to wounding alone (Fig. S5D-E). In contrast, isochorismate synthase 1 (ICS1) homologs with potential roles in inducible SA biosynthesis were unaffected by inceptin treatment (Fig. S5F).

Inceptin also strongly affected expression of defense related transcription factors (TFs) (Fig. S6, Table S7). For specific ethylene response factors (ERFs), inceptin treatment tended to amplify early wound-induced changes. Of the 24 wound regulated ERF transcripts 1 hr after treatment, 4 were further amplified by *Vu*-In treatment, and an additional 2 were *Vu*-In-specific (Table S7). In contrast, inceptin tended to prolong the upregulation of MYC TFs. Transcripts encoding MYC TFs *Vigun03g225300* and *Vigun10g150300* significantly accumulated following wounding alone at 1 hr, but high expression was only prolonged to 6 hr when treated with *Vu*-In (Fig. S6). A smaller proportion of WRKY TF transcripts were affected by *Vu*-In treatment. While 25 WRKY TFs were upregulated by wounding at either 1 hr or 6 hr, only 5 were further affected by inceptin (Table S7). Interestingly, the JAZ family of repressors was very broadly wound and *Vu*-In regulated as demonstrated by the accumulation of 7 of 10 *JAZ* transcripts with *Vu*-In-modulated responses at 6 hr (Fig. S6).

We also identified a targeted subset of gene families likely to be involved in early defense signal transduction that were affected by *Vu*-In (Table S8). Among 32 predicted calmodulin and calmodulin-like (CML) genes, 13 were altered by wounding, but only 2 were further modulated by *Vu*-In. While many of the transcripts encoding 35 predicted calcium-dependent protein kinases (CPK) or 8 predicted mitogen activated protein kinases (MAPK) were significantly accumulated after wounding, none were significantly altered by the further addition of *Vu*-In at either timepoint (Table S8). Taken together, we conclude that addition of *Vu*-In to cowpea wounds leads to targeted effects on signaling pathways and TFs, rather than broadly amplifying entire sets of wound-responsive transcripts.

### *Vu*-In regulated defense-related transcripts encode functional proteins underlaying volatile biosynthesis and direct defense against herbivores

To understand the functional consequences of HAMP-modulated wound responses, we focused on the most highly upregulated genes (log2(FC) > 4) at both 1 hr and 6 hr timepoints, a total of 69 genes (Table 1). Among the most highly *Vu*-In upregulated transcripts at 1 hr were genes encoding a large family of chalcone synthases (CHS), glycosyltransferases, protease inhibitors and cytochrome P450s (CYP) (Table 1). Strongly *Vu*-In upregulated transcripts at 6 hr included genes encoding terpene synthases (TPS) and peroxidases. Transcripts encoding TPS showed especially large, family-wide effects; *Vu*-In positively regulated 8 of 28 cowpea *TPS*, with several *TPS* showing *Vu*-In-specific effects and others a prolonged upregulation from early wound-induced expression (Fig. S7, Table S9). Consistent with late upregulation of specific *TPS* transcripts, *Vu*-In treatment induced the accumulation of 7 terpenes including (3*E*)-4,8-dimethyl-1,3,7-nonatriene [DMNT], *E*-β-farnesene, β-caryophyllene, *E,E*-α-farnesene, linalool, germacrene D and sesquithujene (Fig. 4A). Derived from the tryptophan biosynthetic pathway, indole was also a *Vu*-In elicited volatile. Overall, *Vu*-In elicited leaf volatiles from the current IT97K-499-35 line are highly similar to those previously observed in cowpea variety CB5 (Schmelz *et al*., 2006). To connect genes to functions, we cloned and expressed the most highly upregulated TPS relative to wounding alone, namely *Vigun11g073100*, which displayed a 47-fold *Vu*-In-elicited increase in transcript accumulation at 6 hr (Table S9). Heterologous expression of *Vigun11g073100* in *N. benthamiana* led to production of predominantly germacrene D and 2 comparatively low abundance products, namely ∝-copaene and cubebol (Fig. 4B, Fig. S8). Our results are consistent with *Vigun11g073100* functioning as a germacrene D synthase that contributes to the observed blend of *Vu*-In-elicited cowpea volatiles. Collectively our results indicate that *Vu*-In-modulates wound-induced volatile release in cowpea, corresponding with targeted upregulation of specific *TPS* family members that encode functional sesquiterpene synthases.

**Table 1.**
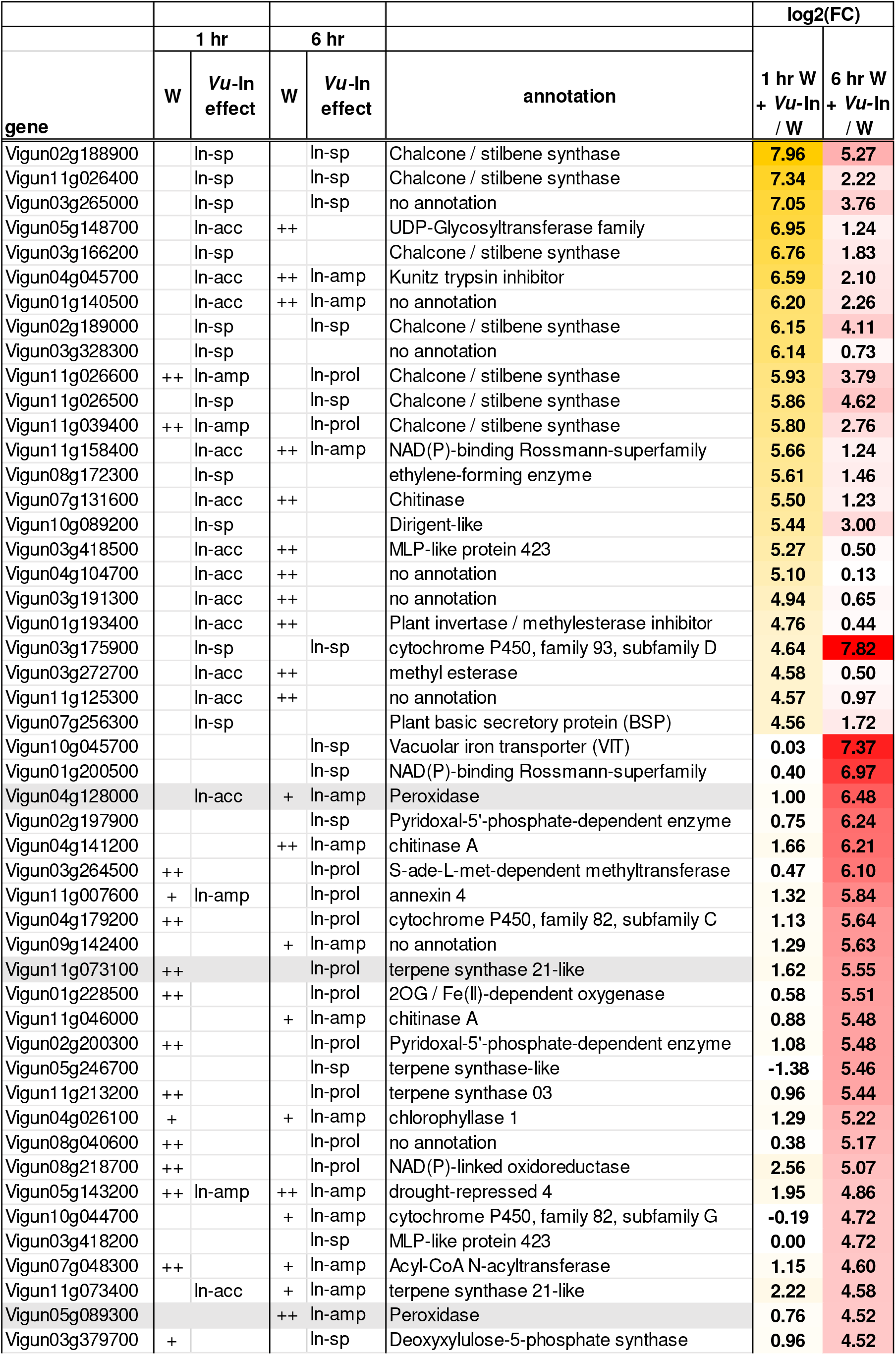
*Vu*-In-elicited transcript accumulation is enriched in defense related gene families. Behavior of significantly accumulating transcripts, log2 Fold Change (FC) > 4 of *Vu*-In vs wounding (W), at either or both of 1 hr and 6 hr timepoints are shown with gene annotations. Under column W, the effect of wounding is indicated for the 1 hr and 6 hr timepoints relative to unwounded tissue collected at 1 hr, where “+” indicates log2(FC) > 1 and Benjamini-Hochberg adjusted p<0.05, “++” indicates log2(FC) > 3. Category of *Vu*-In-induced change is also indicated for both timepoints. *Vu*-In effects on transcript accumulation patterns were categorized as specific (sp), accelerated (acc), amplified (amp), or prolonged (prol). Grey highlighted genes were functionally characterized in this study.

|  | 1 hr |  | 6 hr |  |  | log2(FC) |  |
| --- | --- | --- | --- | --- | --- | --- | --- |
|  | W | Vu-In effect | W | Vu-In effect | annotation | 1 hr W + Vu-In / W | 6 hr W + Vu-In / W |
| gene |  |  |  |  |  |  |  |
| Vigun02g188900 |  | In-sp |  | In-sp | Chalcone / stilbene synthase | 7.96 | 5.27 |
| Vigun11g026400 |  | In-sp |  | In-sp | Chalcone / stilbene synthase | 7.34 | 2.22 |
| Vigun03g265000 |  | In-sp |  | In-sp | no annotation | 7.05 | 3.76 |
| Vigun05g148700 |  | In-acc | ++ |  | UDP-Glycosyltransferase family | 6.95 | 1.24 |
| Vigun03g166200 |  | In-sp |  |  | Chalcone / stilbene synthase | 6.76 | 1.83 |
| Vigun04g045700 |  | In-acc | ++ | In-amp | Kunitz trypsin inhibitor | 6.59 | 2.10 |
| Vigun01g140500 |  | In-acc | ++ | In-amp | no annotation | 6.20 | 2.26 |
| Vigun02g189000 |  | In-sp |  | In-sp | Chalcone / stilbene synthase | 6.15 | 4.11 |
| Vigun03g328300 |  | In-sp |  |  | no annotation | 6.14 | 0.73 |
| Vigun11g026600 | ++ | In-amp |  | In-prol | Chalcone / stilbene synthase | 5.93 | 3.79 |
| Vigun11g026500 |  | In-sp |  | In-sp | Chalcone / stilbene synthase | 5.86 | 4.62 |
| Vigun11g039400 | ++ | In-amp |  | In-prol | Chalcone / stilbene synthase | 5.80 | 2.76 |
| Vigun11g158400 |  | In-acc | ++ | In-amp | NAD(P)-binding Rossmann-superfamily | 5.66 | 1.24 |
| Vigun08g172300 |  | In-sp |  |  | ethylene-forming enzyme | 5.61 | 1.46 |
| Vigun07g131600 |  | In-acc | ++ |  | Chitinase | 5.50 | 1.23 |
| Vigun10g089200 |  | In-sp |  |  | Dirigent-like | 5.44 | 3.00 |
| Vigun03g418500 |  | In-acc | ++ |  | MLP-like protein 423 | 5.27 | 0.50 |
| Vigun04g104700 |  | In-acc | ++ |  | no annotation | 5.10 | 0.13 |
| Vigun03g191300 |  | In-acc | ++ |  | no annotation | 4.94 | 0.65 |
| Vigun01g193400 |  | In-acc | ++ |  | Plant invertase / methylesterase inhibitor | 4.76 | 0.44 |
| Vigun03g175900 |  | In-sp |  | In-sp | cytochrome P450, family 93, subfamily D | 4.64 | 7.82 |
| Vigun03g272700 |  | In-acc | ++ |  | methyl esterase | 4.58 | 0.50 |
| Vigun11g125300 |  | In-acc | ++ |  | no annotation | 4.57 | 0.97 |
| Vigun07g256300 |  | In-sp |  |  | Plant basic secretory protein (BSP) | 4.56 | 1.72 |
| Vigun10g045700 |  |  |  | In-sp | Vacuolar iron transporter (VIT) | 0.03 | 7.37 |
| Vigun01g200500 |  |  |  | In-sp | NAD(P)-binding Rossmann-superfamily | 0.40 | 6.97 |
| Vigun04g128000 |  | In-acc | + | In-amp | Peroxidase | 1.00 | 6.48 |
| Vigun02g197900 |  |  |  | In-sp | Pyridoxal-5'-phosphate-dependent enzyme | 0.75 | 6.24 |
| Vigun04g141200 |  |  | ++ | In-amp | chitinase A | 1.66 | 6.21 |
| Vigun03g264500 | ++ |  |  | In-prol | S-ade-L-met-dependent methyltransferase | 0.47 | 6.10 |
| Vigun11g007600 | + | In-amp |  | In-prol | annexin 4 | 1.32 | 5.84 |
| Vigun04g179200 | ++ |  |  | In-prol | cytochrome P450, family 82, subfamily C | 1.13 | 5.64 |
| Vigun09g142400 |  |  | + | In-amp | no annotation | 1.29 | 5.63 |
| Vigun11g073100 | ++ |  |  | In-prol | terpene synthase 21-like | 1.62 | 5.55 |
| Vigun01g228500 | ++ |  |  | In-prol | 2OG / Fe(II)-dependent oxygenase | 0.58 | 5.51 |
| Vigun11g046000 |  |  | + | In-amp | chitinase A | 0.88 | 5.48 |
| Vigun02g200300 | ++ |  |  | In-prol | Pyridoxal-5'-phosphate-dependent enzyme | 1.08 | 5.48 |
| Vigun05g246700 |  |  |  | In-sp | terpene synthase-like | -1.38 | 5.46 |
| Vigun11g213200 | ++ |  |  | In-prol | terpene synthase 03 | 0.96 | 5.44 |
| Vigun04g026100 | + |  | + | In-amp | chlorophyllase 1 | 1.29 | 5.22 |
| Vigun08g040600 | ++ |  |  | In-prol | no annotation | 0.38 | 5.17 |
| Vigun08g218700 | ++ |  |  | In-prol | NAD(P)-linked oxidoreductase | 2.56 | 5.07 |
| Vigun05g143200 | ++ | In-amp | ++ | In-amp | drought-repressed 4 | 1.95 | 4.86 |
| Vigun10g044700 |  |  | + | In-amp | cytochrome P450, family 82, subfamily G | -0.19 | 4.72 |
| Vigun03g418200 |  |  |  | In-sp | MLP-like protein 423 | 0.00 | 4.72 |
| Vigun07g048300 | ++ |  | + | In-amp | Acyl-CoA N-acyltransferase | 1.15 | 4.60 |
| Vigun11g073400 |  | In-acc | + | In-amp | terpene synthase 21-like | 2.22 | 4.58 |
| Vigun05g089300 |  |  | ++ | In-amp | Peroxidase | 0.76 | 4.52 |
| Vigun03g379700 | + |  |  | In-sp | Deoxyxylulose-5-phosphate synthase | 0.96 | 4.52 |

**Fig. 4.**
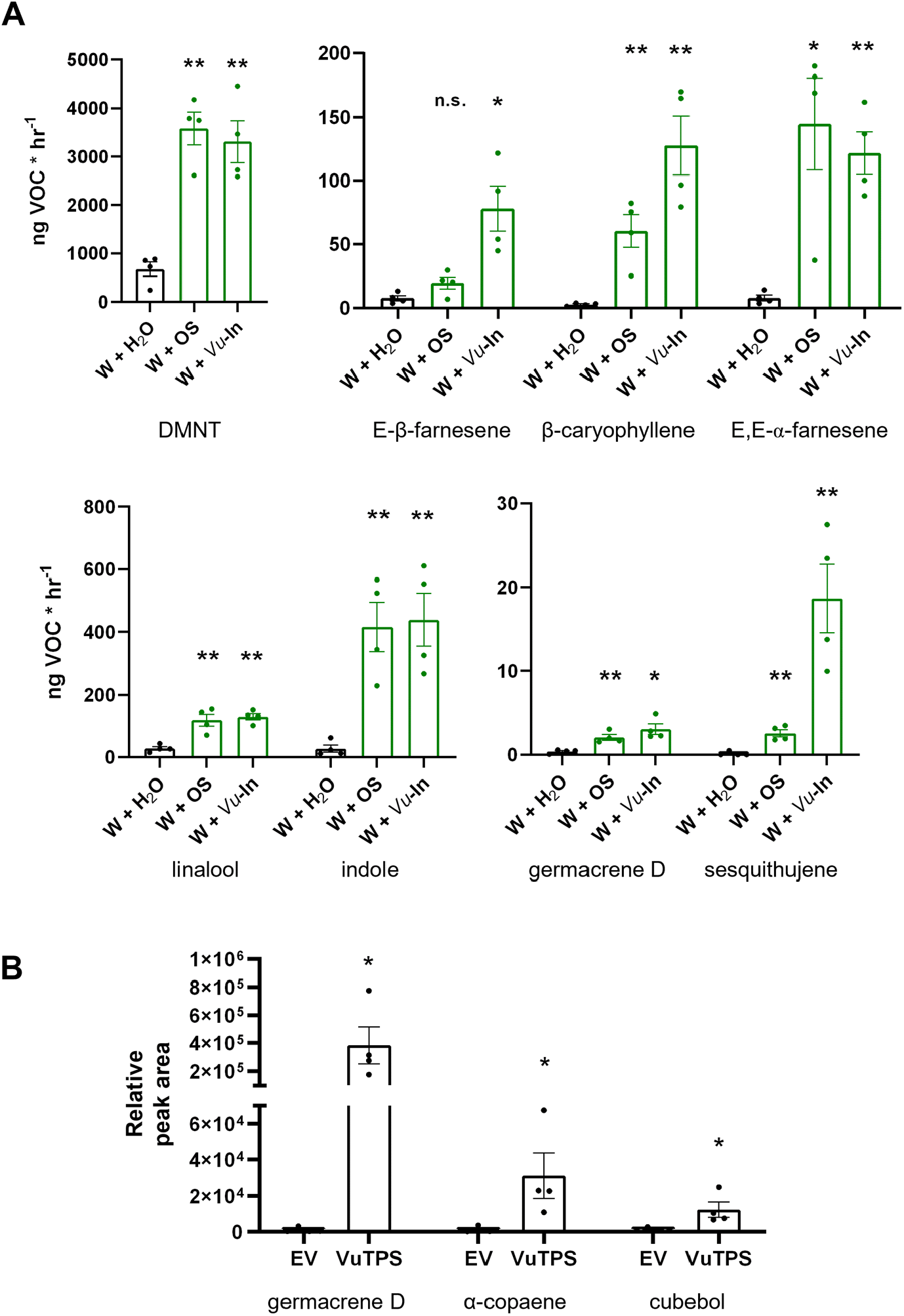
*Vu-*In elicits emission of a blend of cowpea terpene volatiles and accumulation of the transcript *Vigun11g073100* which encodes a functional germacrene D synthase. **A**, Average (N=4, +/-SEM) production of 8 volatile organic compounds (VOC) by cowpea leaves treated with wounding plus either H_2_O (W), *Spodoptera* oral secretions (OS) or *Vu-*In. Asterisk (*) indicates Student’s t-test p<0.05, ** p<0.005. **B**, As an estimate of relative abundance, average (N=4, +/-SEM) peak area (estimate of relative abundance) of volatile sesquiterpene production following transient *Agrobacterium*-mediated heterologous expression of *35S:Vigun11g073100* in *N. benthamiana* leaves. Sesquiterpenes were extracted from whole leaf tissue. Absent from empty vector (EV) controls, *35S:Vigun11g073100* expression results in the predominant production of germacrene D and α-copaene and cubebol. Asterisk (*) indicates 2-tailed Student’s t-test p<0.05

As potential direct defense candidates genes encoding two annotated peroxidases (POX), namely *Vigun04g128000 (VuPOX1)* and *Vigun05g089300 (VuPOX2)*, were also among the most highly induced transcripts at the 6 hr timepoint after wound plus Vu-In treatment (Table 1). *VuPOX1* was also the most *Vu*-In modulated (>24-fold) peroxidase transcript in the genome (Table S10). Consistent with *POX* transcript upregulation generating additional activity, secreted peroxidase activity in excised cowpea leaf discs peripheral to scratch wounded tissue was greater after inceptin treatment relative to wounding alone (Fig. 5A). For both *VuPOX1* and *VuPOX2, Vu*-In treatment amplified wound-induced upregulation relative to undamaged tissue (Fig. S9, Table S10). *VuPOX1* and *VuPOX2* are among several sequence-divergent POX genes induced by inceptin (Fig. S9). Among 97 annotated cowpea POX, inceptin treatment amplified the wound-induction of 8 POX transcripts, representing only half of the 16 total wound-upregulated POX (Fig. S9, Table S10). Our results suggest that a limited number of *Vu*-In responsive *POX* transcripts contribute to *Vu*-In mediated increases in cowpea POX biochemical activity (Fig. 5A).

**Fig. 5.**
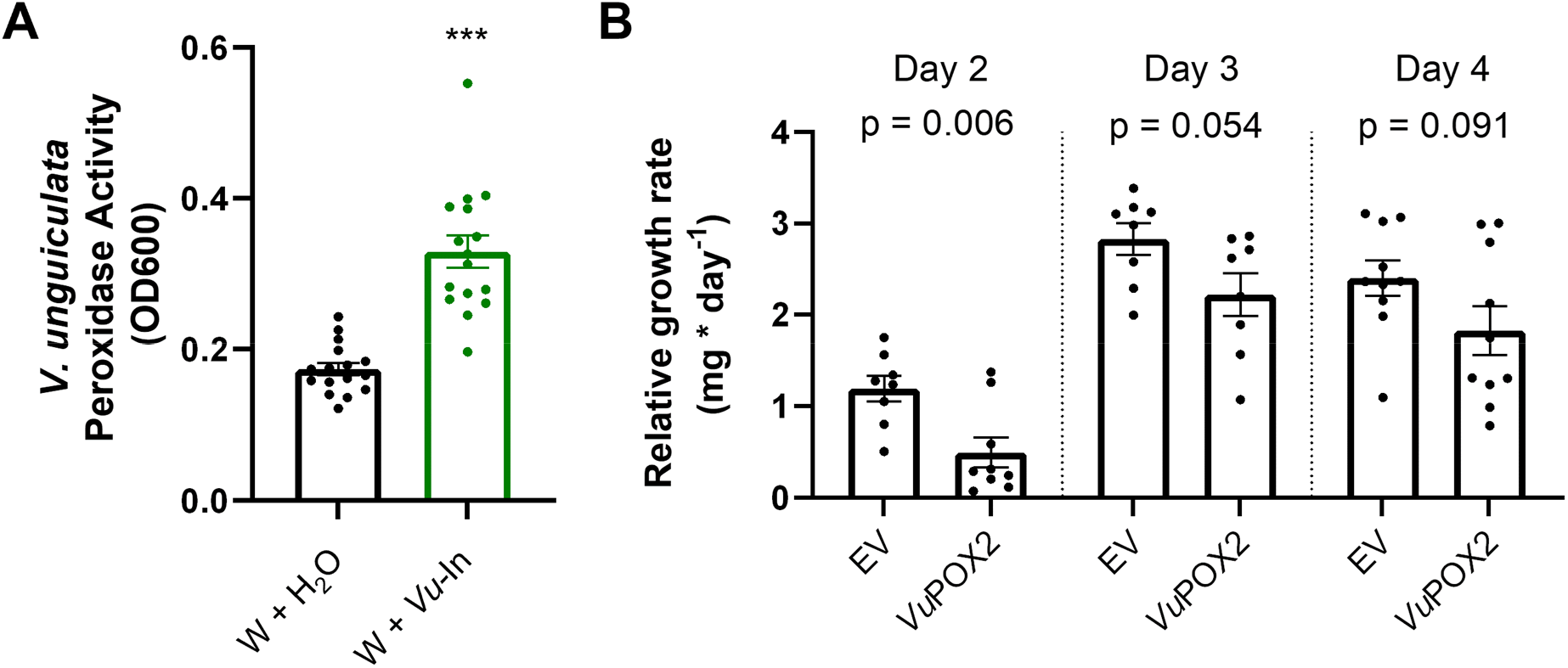
*Vu-*In elicitation increases both peroxidase (POX) activity and *VuPOX* transcript accumulation that is experimentally consistent with a direct defense role against herbivores. **A**, Average (N=15, +/-SEM) peroxidase activity from cowpea in cowpea leaf discs surrounding wound sites treated with either H_2_O or 1 µM *Vu-*In. Asterisk (***) denotes p<0.001, two-tailed Student’s t-test. **B**, Average (n=8-10, +/-SEM) relative growth rate of *Spodoptera exigua* larvae feeding on *N. benthamiana* leaves following the transient *Agrobacterium*-mediated heterologous expression of either an empty vector (EV, pGWB414) or cowpea peroxidase 2 (VuPOX2, *Vigun05g089300*). Results are shown from 3 separate independent experiments with different durations of feeding times, namely 2, 3 or 4 days. P-values of two-tailed Student’s t-test are indicated above VuPOX2 vs EV comparison for each day. Treatment effect of both day and construct are significant by two-way ANOVA for total dataset (F>9.03, p<0.005).

To examine the potential functional role of POX encoded by *Vu*-In regulated *VuPOX1* and *VuPOX2* transcripts contributing to antiherbivore direct defenses, we conducted heterologous protein expression assays to assess changes in insect growth. We cloned and expressed *VuPOX1* and *VuPOX2* transiently in N. benthamiana leaf tissue. While *VuPOX1* did not reduce beet armyworm (*Spodoptera exigua*) growth (Fig. S10), VuPOX2 significantly reduced *S. exigua* larval growth rates in separate repeated trials (Fig. 6B). Our results support *Vu*-In-induced POX activity as one of the layers of direct cowpea defenses against caterpillar herbivores.

## V. Discussion

### The model HAMP inceptin specifically modulates the cowpea wound response

Plant perception of HAMPs significantly drives the activation of antiherbivore defenses; however, precise molecular interactions and layered contributions onto wound-induced responses remain less clear. Our current effort details transcriptomic analyses of wounding in the absence and presence of the potent legume-specific HAMP, inceptin. While previous efforts have demonstrated a role for crude OS in amplifying wound responses or activating specific outputs (Bricchi *et al*., 2010), our work uniquely focuses on a single defined HAMP with established receptor-ligand immune recognition (Steinbrenner *et al*., 2020). Since *Vu-*In elicitation occurs via a dedicated pattern recognition receptor, any changes can be interpreted as true HAMP-induced responses, rather than the output of a combination of elicitors and suppressors. Thus the hundreds of specific transcriptional targets represent diverse processes where wound-induced changes can be *Vu*-In amplified, *Vu*-In accelerated, or occur specifically in the context of HAMP elicitation.

Previous transcriptomic analyses using defined HAMPs did not directly compare patterns to the plant wound response. For example, Zhou *et al*. (2016) compared wounded tobacco plants treated with the HAMP 18:3-Gln to wounded, water-treated controls and detected up to ∼1800 upregulated genes across three early timepoints; however, the baseline expression levels in unwounded plants was not measured. Other studies have measured responses to total OS or live herbivores and measured similarly large sets of inducible transcripts (Appel *et al*., 2014; Gilardoni *et al*., 2011; Consales *et al*., 2012). Our current study allows a direct comparison with unwounded plant transcriptional behavior at 1 hr, and an estimated effect of wounding at 6 hr relative to the 1 hr (ZT4) samples, revealing distinct patterns relative to wounding alone (Fig. 3, Fig. S4). We thus build upon previous efforts by using a known HAMP to define specific transcriptional changes within the context of the plant wound response (Reymond *et al*., 2000; Major and Constabel, 2006). Several caveats still apply: first, artificial scratch-wounding may not capture effects of repeated bouts of wounding during live herbivory, or reflect the dynamics of *Vu-*In exposure to relevant cell populations (Bricchi *et al*., 2010; Farmer *et al*., 2020). Second, the pool of cells collected for analyses cannot reflect spatial heterogeneity around a wound site, including the role of signaling components in vascular cells (Nguyen *et al*., 2018). Our study nonetheless reflects one possible set of wound-HAMP interactions with gene-level resolution.

As a global trend, we collectively observed a greater number of upregulated genes than downregulated genes (Fig. 1B). By including unwounded control tissue at 1 hr and measuring both early (1 hr) and late (6 hr) timepoints for wounding +/-*Vu*-In, we were able to categorize genes specifically modified by inceptin but not by wounding alone at 1 hr and estimate wound effects at 6 hr. Overall, at either 1 hr and 6 hr timepoints, *Vu*-In amplified the wound effect for 541 genes, led to specific changes for 715 genes, and either prolonged or accelerated the wound-induced changes for 328 genes (Fig. 3, Tables S2-S3). Thus, the legume HAMP *Vu*-In does amplify select wound responses; however, the category of *Vu*-In-amplified transcripts represents only a third of all *Vu*-In-induced transcriptional changes. Within specific gene families, where *Vu*-In amplified transcript accumulation for only a few genes among a broader set of wound-induced responses (Figs. S5, S6). Recognition of the HAMP *Vu*-In has highly targeted effects that contribute to the total cowpea response to herbivory that includes wounding.

Surprisingly, wound plus inceptin treatment reversed the transcriptional behavior of many genes relative to wounding alone. For example, at the 1 hr timepoint, *Vu*-In interacted with early wound-repression exclusively as a positive regulator; no wound-downregulated genes were further downregulated by addition of *Vu*-In, while 53 genes were significantly reversed in a positive direction by *Vu*-In. Reversal of wound response was common for wound-upregulated transcript accumulation as well. At both 1 hr and 6 hr, a total of 201 genes had significant sign reversals relative to the effect of wounding alone (Table S4, S5). While 6 hr changes relative to undamaged tissue at 1 hr should be interpreted with caution, wound-reversed changes at 1 hr reflect a comparison controlled for circadian effects. Previous studies in different model systems have noted that OS treatment can suppress defense-related transcription factors, and this was proposed to result from active signaling suppression by insect effectors (Consales *et al*., 2012; Appel *et al*., 2014; Chen *et al*., 2019). By using a single defined HAMP we also observe suppression signatures suggesting an alternative interpretation where plants actively modulate or suppress wound-induced transcription after perception of a potent HAMP in the absence of additional negative effectors. Transcripts that are reversed by *Vu*-In treatments encode enzymes, transporters, and proteins of unknown function that exist as candidates for highly tuned defense outputs to chewing herbivory.

### Damage gating of HAMP and PAMP responses in plants

In many plants the full set of herbivore-inducible defense responses requires signals beyond wounding alone. For example, early analyses of *Vu*-In outputs in cowpea are consistent with a HAMP requirement for volatile emission, such as (*E*)-4,8-dimethyl-1,3,7-nonatriene (DMNT), and multiple phytohormone changes (Schmelz *et al*., 2006). At the transcriptome wide level, our current analyses are consistent with a requirement of HAMP perception for a large spectrum of defensive outputs. Importantly, the current transcriptomic analyses also enable comparisons to PAMP- and DAMP-induced changes in other model systems. Denoux *et al*. (2008) compared Arabidopsis responses to flg22 and oligogalacturonides, and observed distinct early and late transcriptional changes from 1 to 12 hr after treatment. Similar patterns have been observed in other flg22-induced transcriptomic analyses, generally transitioning from signaling factors to classic hormonal and defensive outputs such as pathogenesis-related (PR) transcripts (Hillmer *et al*., 2017; Göhre *et al*., 2012; Roux *et al*., 2011; Bjornson *et al*., 2021). Our analysis of *Vu-*In-induced changes reveals a similar pattern; distinct sets of genes are modified by inceptin treatment at 1 hr and 6 hr, in broadly different sets of gene families (Fig. 2B, Table S2-S3).

The observed HAMP responses are also consistent with a damage requirement for full elicitor responsiveness, which is a growing theme in plant immunity to microbial pathogens. In Arabidopsis roots, mechanical damage gates plant responsiveness to the flg22 PAMP through upregulation of the corresponding flg22 receptor FLS2 in neighboring cells (Zhou *et al*., 2020), potentially minimizing immune activation by beneficial microbes. In interactions with chewing herbivores, plant exposure to *Vu*-In should similarly not occur without also imparting mechanical wounding. Inceptin responses in cowpea are mediated by the recently described LRR-RLP INR (Steinbrenner *et al*., 2020), and while the *INR* gene itself is not significantly modified by wounding or inceptin, other DAMP receptors or DAMP-induced transcriptional changes may functionally interact with HAMP perception at early signaling steps (Huffaker *et al*., 2013; Shinya *et al*., 2018). Further investigation of interfaces between INR and DAMP signaling will reveal the molecular basis for HAMP-amplified and HAMP-specific transcriptional changes.

### Targeted subsets of defense-related gene families are inceptin regulated

In less commonly targeted model organisms, such as cowpea, pipelines for assigning gene annotations may imperfectly identify putative functions. To reduce this problem, one approach is to frame gene family members in phylogenetic context with functionally characterized orthologs. Towards this goal, we placed *Vu*-In regulated gene family members of ethylene / JA biosynthesis, response, and signaling factors in the context of sequence similarity to defined Arabidopsis genes (Figs. S5-S6, Table S7-S8). For example, *Vu-*In amplified the wound-induced expression of *Vigun02g178400*, an ortholog of ACS6 in Arabidopsis, while other wound-induced ACS genes were not further affected by *Vu-*In (Fig. S5). The timing of transcriptional activation of hormone biosynthetic machinery roughly matches the timeline of induced hormone responses previously observed in cowpea (Schmelz *et al*., 2007).

*Vu-*In-specific effects were especially notable for a subset of JAZ transcripts, *Vigun03g013500* and *Vigun02g021400*, orthologs of Arabidopsis JAZ9. JAZ9 has been specifically implicated in regulation of the ERF1/ORA59-mediated branch of JA signaling (Takaoka *et al*., 2018), and late upregulation of JAZ9 homologs by *Vu*-In may prevent long-term accumulation of highly specific outputs downstream of ERF but not MYC TFs. Consistent with this model, close homologs of ERF1 and ORA59, namely *Vigun07g099700* and *Vigun07g178200*, were among the few *Vu-*In upregulated ERFs at 1 hr, but their expression levels generally fall by 6 hr (Table S7). HAMP-induced changes were also targeted in CML, CPK, and MAPK pathways; however, only 2 of 20 wound-induced transcripts were further modulated by inceptin (Table S8). Further functional insights into regulation of HAMP-induced signaling factors will require additional reverse genetic tools in cowpea, for example TILLING populations or reliable transformation to introduce CRISPR/Cas mediated mutations.

### Relationship to Herbivore-induced Terpenes

Strong *Vu*-In-specific changes relating to defense outputs were predominated by specific biosynthetic pathways unique to either the early or late timepoints. For example, *Vu*-In treatment elicited the specific accumulation of a large array of transcripts at 1 hr encoding predicted chalcone synthases (Fig 4). This parallels related legume models where *Spodoptera litura* gut contents strongly elicited the biosynthesis of isoflavones (Nakata *et al*., 2016). Similarly *Vu*-In-amplified and *Vu*-In-prolonged expression of terpene synthase (TPS) transcripts at 6 hr (Fig. 4) matches earlier predictions based on *Spodoptera* OS and *Vu*-In-elicited volatiles production in cowpea leaves (Schmelz *et al*., 2006). Collectively the strong elicitation of CHS and TPS transcripts tracks the common co-occurrence of flavonoids and terpenoids (Ding *et al*., 2020; Lange, 2015). Consistent with the present transcriptomic results, cowpea plants scratch-wounded and treated with *Vu*-In produced a variety of monoterpenes, homoterpenes and sesquiterpenes as well as greater peroxidase activity than mock treated plants (Fig. 5A, 6A).

We functionally characterized the most strongly upregulated TPS transcript, *Vigun11g073100*, at the late 6 hr timepoint, where *Vu*-In strongly prolonged wound-induced early expression (Table 1, Fig. S7). Following *Vigun11g073100* expression in *N. benthamiana*, we detected de novo production of germacrene D and two related minor sesquiterpene products. Interestingly, germacrene D is detectable following *Vu*-In elicitation but is a comparatively low abundance sesquiterpene emitted by cowpea leaves (Fig. 5B). It is likely that other synchronously accumulating TPS transcripts (Fig. 4) have a larger contribution to the measured blend of *Vu*-In elicited terpenes in cowpea. Co-regulated cytochrome P450s could also drive the production of non-volatile terpene defenses and also contribute to the regulation of volatile emission in complex ways (Liu *et al*., 2015). *Vigun11g073100* belongs to a highly legume specific clade of TPS and contains features of Type I monofunctional TPS including alpha, beta, but not gamma helical domains and intact Type I DDxxD motif, consistent with characterized enzymes catalyzing sesquiterpene biosynthesis (Zhou and Pichersky, 2020). Other TPS with similar *Vu*-In mediated expression patterns are phylogenetically distant from *Vigun11g073100* (Fig. S7) and may contribute separate functions, consistent with the variety of inceptin-induced terpenes observed in cowpea (Fig. 5B). Importantly, caterpillar-elicited volatiles in cowpea leaves include many minor and unidentified compounds that produce significant electroantennogram responses in diverse parasitioid wasp species (Gouinguené *et al*., 2005). Highly abundant HAMP elicited volatiles serve as useful research biomarkers but need not serve as the most important cues mediating indirect defense responses.

### Relationship to Defense-Related Peroxidase Activities

As candidates for direct defenses, *Vu*-In also strongly influenced the accumulation of specific peroxidase transcripts, greatly amplifying both measurable enzymatic activity of secreted leaf peroxidases and the wound-mediated transcription of 6 members of the large gene family (Fig. 6A,B). We functionally characterized the two most strongly *Vu*-In-regulated *POX* transcripts, *Vigun04g128000* and *Vigun05g089300* (Fig. 6B). In heterologous expression assays using *N. benthamiana*, only *Vigun05g089300* consistently suppressed the relative growth rate of attacking *S. exigua* larvae (Fig. 6C, Fig. S10). Collectively our results link specific *Vu*-In regulated transcript levels to cowpea enzymatic activities and antiherbivore defense-related outputs when examined in heterologous systems. Both *VuPOX1* and *VuPOX2* encode Class III enzymes, distinct from Class I Ascorbate Peroxidases with potential anti-herbivore activity via decreasing ascorbate availability in the insect gut (Felton and Summers, 1993). Class III peroxidases instead form a large gene family in plants with diverse substrates and functional roles in scavenging/generating reactive oxygen species and in cell wall fortification (Shigeto and Tsutsumi, 2016). For example, silencing Arabidopsis class III peroxidase Prx34 impaired both reactive oxygen species production and pathogen defense (Daudi *et al*., 2012). The specific mechanism for defensive activity of *VuPOX2* is not known and may reflect substrate specificity relevant to digestive processes in chewing herbivores. It will also be interesting to compare HAMP-inducible *POX* transcripts to alternative bacterially-responsive transcripts (Wang *et al*., 2018). It is possible that regulatory or functional specificity of pathogen-induced peroxidases amplifies alternative branches of the plant immune system, antagonizing defense against chewing herbivores. Given that peroxidases play an active role in plant redox immune signaling, future characterization of PAMP vs. HAMP induced POX gene family members may provide clues for specific functions.

### Model for coordination of DAMP and HAMP signaling

Our work highlights HAMP specific defense contributions to wound-induced transcriptional reprogramming in a legume crop plant. We present a model where a combination of wound- and receptor-mediated DAMP signaling underlies most transcriptional changes at a local cowpea wound site, yet the addition of defined HAMPs as a further immunogenic input further amplifies and modulates the wound response resulting examples of response specificity. *Vu*-In is recognized by the recently described inceptin receptor, INR, providing a molecular probe to assess herbivore specific plant responses. Our current effort provides a foundation of HAMP-specific responses in legumes necessary to provide answers to a fundamental question namely “How can innate plant immune responses be specifically tuned for anti-herbivore defenses?” In the future, the molecular interactions of inceptin-INR signaling and outputs examined alongside classical wound and PAMP signaling pathways can provide key insights to understand how plants discriminate between herbivores and pathogens after the perception of key molecular patterns.

## VI. Experimental Procedures

### Plant Growth and Treatments

Genome-sequenced cowpea variety IT97K-499-35 was used for all experiments. Cowpea seeds were germinated and grown in a walk-in growth chamber (22°C, 12/12 day night cycle) in 3-inch pots with Berger Mix 2 (Berger, Saint-Modeste, Quebec, Canada). For transcriptomic analyses, n=4 plants were harvested as biological replicates for each of 5 treatments: 1) 1 hr unwounded, 2) 1 hr wounded + H_2_O, 3) 1 hr wounded + *Vu*-In, 4) 6 hr wounded + H_2_O, 5) 6 hr wounded + *Vu*-In. Trifoliate leaves of 3-week-old plants were initially treated at 11 AM (ZT4). For wounding, the adaxial sides of new fully expanded leaves were superficially scratched with a razor in three areas, removing 2.5% of the waxy cuticle. The wound sites included the central leaf tip spanning both sides of the midrib and two midbasal sections on opposite sides of the midrib. Using a pipette, 10 μL of an aqueous solution containing either H_2_O only or 1 μM *Vu*-In were immediately applied and dispersed over the damage sites. Leaf punches (0.26 mg, 2 cm diameter) containing the entire wound plus peripheral tissue were collected at 1 hr and 6 hr timepoints as described above.

For the analyses of *Vu-*In elicited cowpea volatiles, trifoliate leaves of 3-week-old greenhouse grown cowpea plants were used. Plants were scratch-wounded and treated with H_2_O, cowpea-fed *Spodoptera frugiperda* oral secretions collected as described (Schmelz *et al*., 2006), or *Vu*-In as described above. To ensure strong plant responses for the analyses of elicited volatile emission, leaf treatments were repeated 3 times as follows: 7:30 AM and 6:30 PM on day one and again 7:30 AM the following morning on day 2. One hr later, starting at 8:30 AM a 1 h volatile collection from individual excised cowpea leaves was performed following (Schmelz *et al*., 2001).

### Transcriptomic Analysis

Total RNA was extracted using Sigma Plant RNA Kit (Sigma-Aldrich, St. Louis, MO). 3’ TruSeq Library construction was performed on total RNA at Cornell Institute of Biotechnology. Reads were trimmed with Trimmomatic (options LEADING:3 TRAILING:3 SLIDINGWINDOW:4:15 MINLEN:36 HEADCROP:12) and were mapped to the cowpea reference genome (Lonardi *et al*., 2019) using Hisat2 (Min intron length 60, Max intron length 6000) (Kim *et al*., 2019). Transcript counts for individual genes were quantified using HTSeq using default settings (Anders *et al*., 2015). Count data was analyzed for differential expression using DESeq2 (Love *et al*., 2014). Normalized count data is reported in all figures. Differential expression thresholds were log2(fold-change) > 1 and Benjamini-Hochberg adjusted p < 0.05 unless indicated.

### Phylogenetic methods

Cowpea transcripts were compared to Arabidopsis homologs using a custom pipeline. The protein sequence of a single gene family member was used as a query for TBLASTN search of cDNA annotations of both cowpea (v.1.1) and Arabidopsis (TAIR10) genomes. The full protein sequence of all blast hits was compiled and aligned using ClustalOmega (Sievers and Higgins, 2014). Phylogenetic trees were generated using FastTree (Price *et al*., 2010). Trees were visualized with associated transcriptomic data using ggTree (Yu *et al*., 2017).

### Molecular cloning and TPS heterologous expression

Cowpea TPS and POX genes were amplified from cDNA and cloned into a modified pENTR using primers in Table S11. For transient expression of the cowpea TPS in *Nicotiana benthamiana*, a pEarleyGate100 construct (Earley *et al*., 2006) containing the full *Vigun11g073100* ORF was constructed using Gateway cloning and electroporated into *Agrobacterium tumefaciens* strain GV3101. The *Agrobacterium* strain, along with a strain encoding a truncated cytosolic *Euphorbia lathyrism* 3-hydroxy-3-methylglutaryl-coenzyme A reductase (HMGR; EIHMGR, JQ693150.1) was used to increase sesquiterpene accumulation *in planta*. A separate *Agrobacterium* strain encoding the P19 protein to suppress gene silencing in *N. benthamiana* was also included. All *Agrobacterium* strains were cultured overnight and subsequently diluted to a final OD_600_ of 0.8 in 10mM MES pH 5.6, 10mM MgCl_2_. Equal volumes of each culture were mixed, then infiltrated into the youngest set of fully expanded leaves on 30-day-old *N. benthamiana* plants via needleless syringe. Leaf tissues were frozen in liquid N_2_ five days post-infiltration then stored at -80°C for further analysis.

*N. benthamiana* leaf tissue sesquiterpene pools were extracted using a modified vapor-phase extraction (VPE) protocol (Schmelz *et al*., 2004), substituting hexanes for MeCl_2_ for all extraction steps. GC/MS analysis was conducted as described below for cowpea leaf volatiles. 2*sqrt(3/8+X) transformation was applied to all data prior to analysis to account for elevated variation associated with larger mean values.

### Analyses of *Vu*-In-elicited cowpea leaf volatiles

Cowpea leaf volatiles were obtained by passing air over the leaves in a glass cylinder and trapping compounds on inert filters containing 50 mg of HayeSep Q (80- to 100-μm mesh) polymer adsorbent (Sigma-Aldrich) for 1 hr. Description and construction of the volatile collection filters follow from Schmelz et al. (2004). Volatile analytes were eluted from the filters with 150 mL of 1:1 hexane:ethyl acetate containing 2500 ng of nonyl acetate as an internal standard. Gas chromatography (GC) / mass spectrometry (MS) (GC/MS) analyses utilized an Agilent 6890 series GC coupled to an Agilent 5973 mass selective detector (interface temperature, 250°C; mass temperature, 150 °C; source temperature, 230°C; electron energy, 70 eV). The GC was operated with a DB-35MS column (Agilent; 30 m × 250 μm × 0.25 μm film). Cowpea volatiles were introduced with a 1 mL splitless injection and an initial oven temperature of 45 °C. The temperature was held for 2.25 min, then increased to 300 °C with a gradient of 20°C min^−1^ and held at 300°C for 5 min. GC/MS-based estimated quantities of the dominant detected analytes by order of retention time included (3*E*)-4,8-dimethyl-1,3,7-nonatriene [DMNT; retention time (RT) 7.67 min, *m*/*z* 69], linalool (RT 7.71 min, *m*/*z* 93), internal standard nonyl acetate (RT 9.25 min, *m*/*z* 98), sesquithujene (RT 9.70 min, *m*/*z* 93), *E*-β-farnesene (RT 10.16 min, *m*/*z* 93), β-caryophyllene (RT 10.20 min, *m*/*z* 93), indole (RT 10.26 min, *m*/*z* 117), *E,E*-ɑ-farnesene (RT 10.56 min, *m*/*z* 93), and germacrene D (RT 10.69 min, *m*/*z* 161). Volatile compounds were identified by comparison of retention times with authentic standards and by comparison of mass spectra with the Adams, Wiley and National Institute of Standards and Technology libraries. *Vu*-In elicited cowpea leaf volatiles detected in the current study with IT97K-499-35 are highly similar to those previously reported in the commercial line California Blackeye no. 5 (Schmelz *et al*., 2006).

### Peroxidase assays

3-week old cowpea plants were exposed to HAMP elicitation by scratch wounding and *Vu*-In peptide treatment in transcriptomic experiments. Two hours after the treatment, 4mm diameter leaf discs were punched from the periphery of the scratch wound and washed in 1x Murashige-Skoog (MS) solution on a rotary shaker for 1 hr. Leaf discs were then individually placed in a 96-well Clear Flat Bottom Plate and incubated in 50 μL of 1x MS for 20 hours to accumulate secreted peroxidases. After 20 hours, the leaf discs were removed from the 1x MS media and peroxidase activity was measured as described in Mott *et al*. (2018). Briefly, 50 μL of 1 mg/mL 5-aminosalicylic acid solution (pH 6.0 w/ 0.01% hydrogen peroxide) was added to each well. After two minutes, 20 μL of 2M NaOH solution was added to each well to quench the reaction. OD600 absorbance was measured immediately.

### POX heterologous expression and Herbivory assays

POX genes were cloned into pGWB414(Nakagawa *et al*., 2007) with C-terminal 3xHA tag. Three week old *N. benthamiana* plants were infiltrated with *Agrobacterium tumefaciens* (OD_600_ = 0.45) containing empty vector or 35S-driven cowpea POX genes *Vigun04g128000* (*VuPOX1*) or *Vigun05g089300* (*VuPOX2*). Eight leaf punches (#7 cork borer, ∼0.26 mg of tissue per disc) were extracted and placed in petri dishes with individually pre-weighed single 2^nd^ instar beet armyworm larvae in a petri dish, humidified with 0.5 mL H_2_O soaked into a Kimwipe. Larvae were kept in dark incubator for 2-4 days at 28 C and relative growth rate was calculated as weight gained divided by number of days (Waldbauer, 1968).

## VII. Accession Numbers

Transcriptomic data are available in NCBI SRA (BioProject PRJNA758058).

## VIII. Acknowledgements

We acknowledge Alisa Huffaker, Elly Poretsky, and other members of the Huffaker and Schmelz labs for valuable discussion. UCSD STARS (Summer Training Academy for Research Success) provided funding and mentorship structures for NJH. A.D.S. was partially supported by E.A.S. University of California San Diego (UC San Diego) startup funds, the HHMI Postdoctoral Fellowship through the Life Sciences Research Foundation, the University of California President’s Postdoctoral Fellowship Program, and startup funding as a Washington Research Foundation Distinguished Investigator. E.A.S. acknowledges partial support from the UC San Diego startup funds and USDA NIFA AFRI #2018-67013-28125.

## IX. Supporting Information

**Table S1**, DESeq2 expression comparisons

**Table S2**, *Vu*-In-upregulated and downregulated genes at 1 hr

**Table S3**, *Vu*-In-upregulated and downregulated genes at 6 hr

**Table S4**, *Vu*-In reversed genes at 1 hr relative to wounding (W) vs undamaged tissue

**Table S5**, *Vu*-In reversed genes at 6 hr relative to wounding (W) vs undamaged tissue

**Table S6**, Gene Ontology (GO) categories for biological processes of *Vu*-In-regulated genes

**Table S7**, *Vu*-In and wound-induced responses of A, ethylene response factor (ERF) and B, WRKY family transcription factors

**Table S8**, *Vu*-In and wound-induced responses of A, calmodulin and calmodulin-like, B, calcium-dependent protein kinase (CPK), and C, mitogen-activated protein kinase (MAPK) gene family members

**Table S9**, Transcriptional *Vu*-In- and wound-induced (W) responses of terpene synthase (TPS) gene family members

**Table S10**, Transcriptional *Vu*-In- and wound-induced (W) responses of peroxidase (POX) gene family members

**Table S11**, Primers used in the study

**Fig. S1**, Heatmap of clustered log2(fold-change) measures by gene

**Fig. S2**, Alluvial diagram of transcriptional behavior of differentially expressed genes (DEG) after wounding (W) or W + *Vu-*In treatment relative to undamaged tissue (1 hr)

**Fig. S3**, Summary of filtering criteria for categorization of *Vu-*In-induced differential expression

**Fig. S4**, Transcriptional regulation of *Vu-*In-downregulated genes at 1 hr and 6 hr relative to wounding alone

**Fig. S5**, Wound- and *Vu-*In-induced changes in gene expression for defense hormone biosynthetic gene families

**Fig. S6**, Wound- and *Vu-*In-induced changes in gene expression for gene families involved in jasmonate-related transcriptional regulation

**Fig. S7**, Wound- and *Vu*-In-induced changes in gene expression for terpene synthases (TPS)

**Fig. S8**, TIC trace from GC/MS analysis of derivatized *N. benthamiana* volatile pools

**Fig. S9**, Wound- and *Vu-*In-induced changes in gene expression for peroxidases (POX)

**Fig. S10**, Expression of *VuPOX2* but not *VuPOX1* in *N. benthamiana* lowered *S. exigua* growth rates

## XIII. Supplementary Figures

**Fig. S1.**
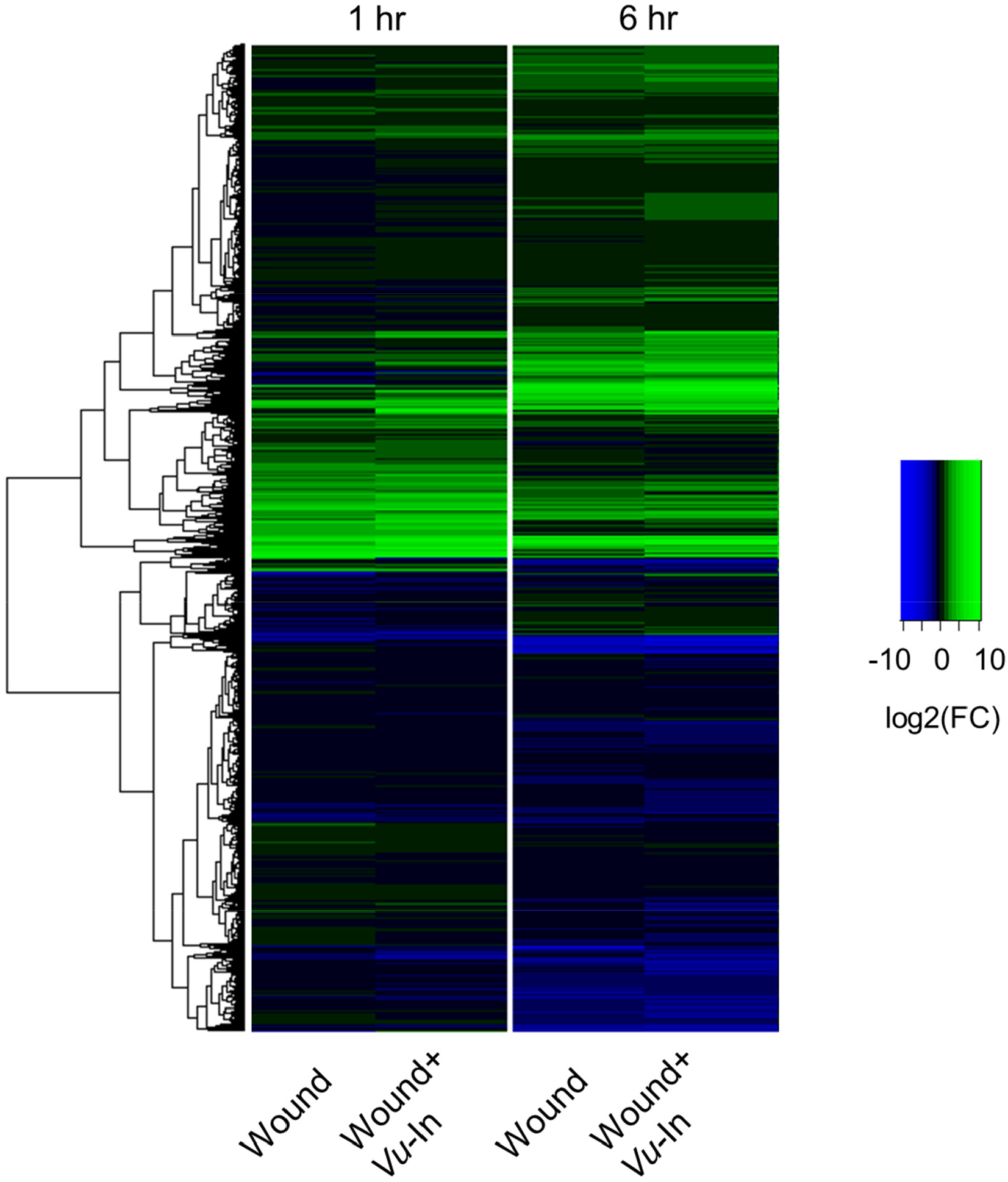
Heatmap of log2(fold-change) measures by gene. Color scale indicates log2(FC) for each treatment-timepoint combination relative to unwounded plants collected at 1 hr. Dendrogram indicates hierarchal clustering using matrix of log2(FC) values for 29,773 genes. Green color indicates magnitude of upregulation and blue color indicates magnitude of downregulation relative to unwounded sample according to legend.

**Fig. S2.**
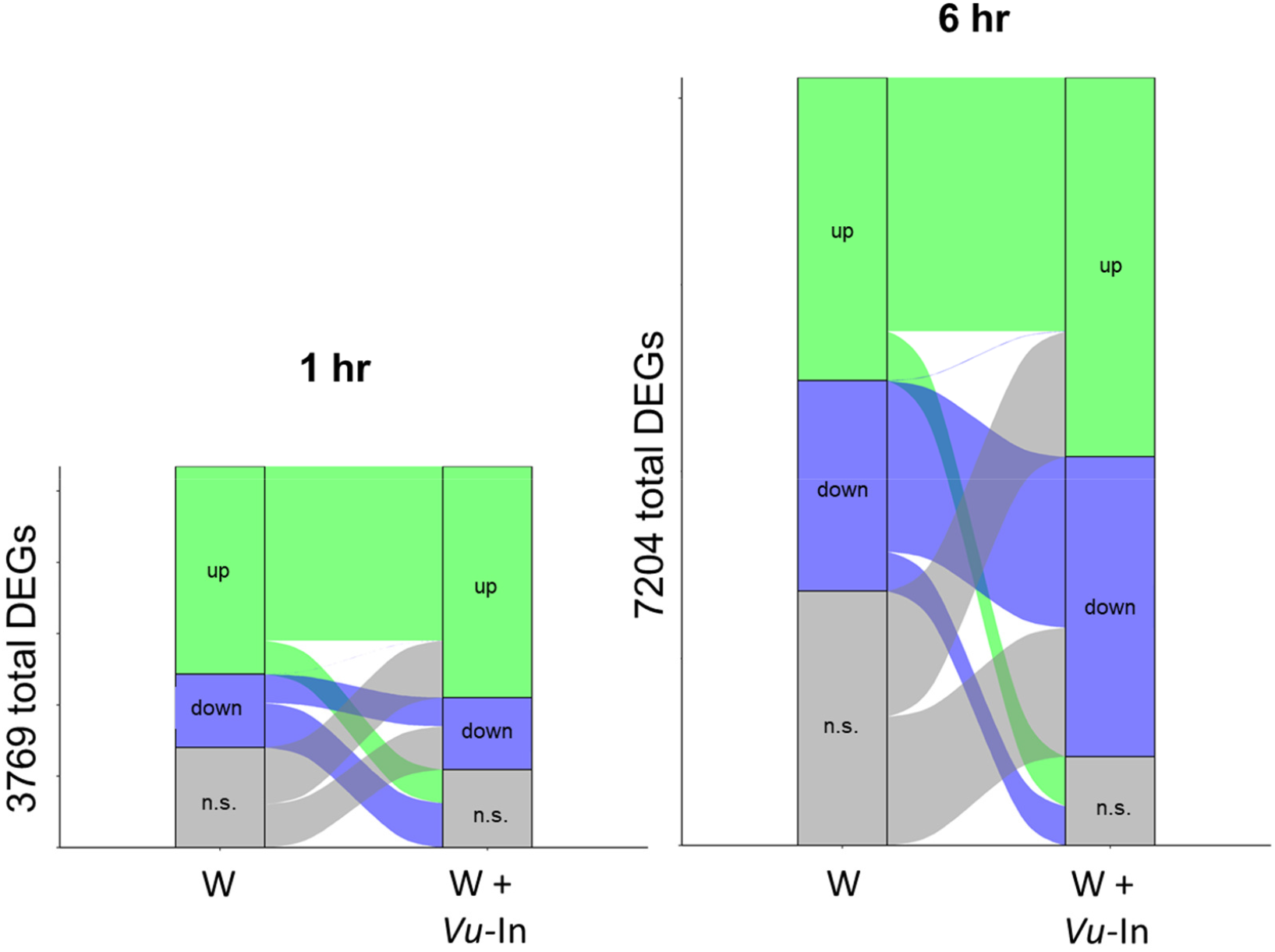
Alluvial diagram of transcriptional behavior of differentially expressed genes (DEG) after wounding (W) or W + *Vu-*In treatment relative to undamaged tissue (1 hr). All DEGs (log2FC > 1, Benjamini-Hochberg adjusted p<0.05) relative to undamaged tissue are included for each timepoint. Genes are binned according to significant upregulation (“up”) or downregulation (“down”) relative to undamaged leaf tissue. n.s., not significantly different from undamaged treatment. Bar height indicates proportional number of genes in each category. Sets of genes with different or similar categorizations in different treatments are connected through alluvial flows between W and W+*Vu*-In treatments.

**Fig. S3.**
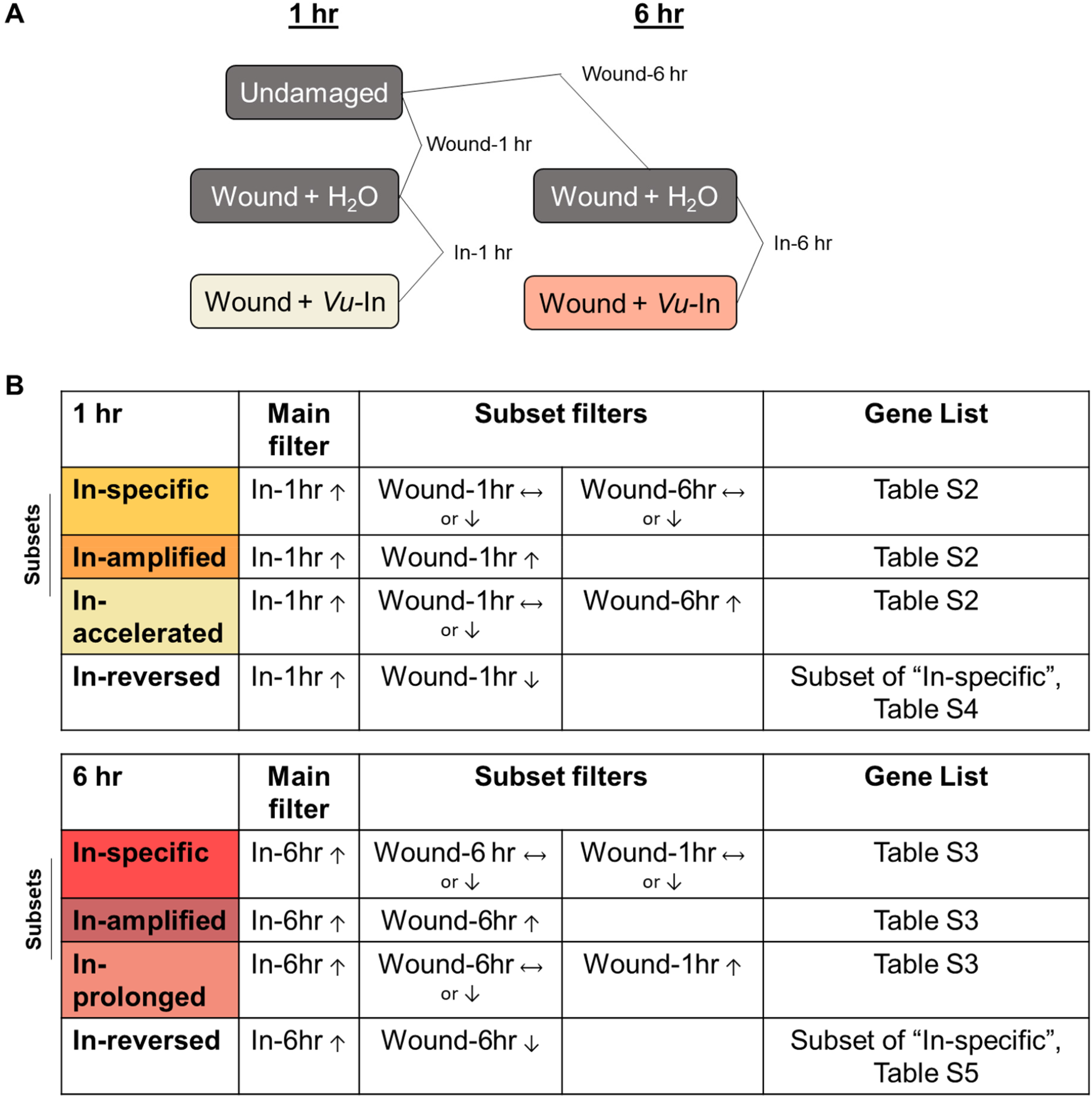
Summary of filtering criteria for categorization of *Vu-*In-induced differential expression. A, Timepoints and treatments harvested for RNA-seq analysis. Lines indicate comparisons of gene expression performed in DESeq2 used to filter genes into sets according to significant differential expression (log2FC > 1, Benjamini-Hochberg adjusted p<0.05) B, Subset filtering criteria for inceptin-upregulation gene expression at 1 hr and 6 hr, based on behavior of the same genes in response to wounding at the same timepoints (subset filters). Wound-induced behavior is relative to 1 hr timepoint as in panel A. Arrows indicate that a given treatment induces upregulation, downregulation, or no significant change (horizontal arrows). Similar criteria were used for downregulation in Fig. S4. Supplemental tables are indicated with gene lists and quantitative behavior of specific subsets.

**Fig. S4.**
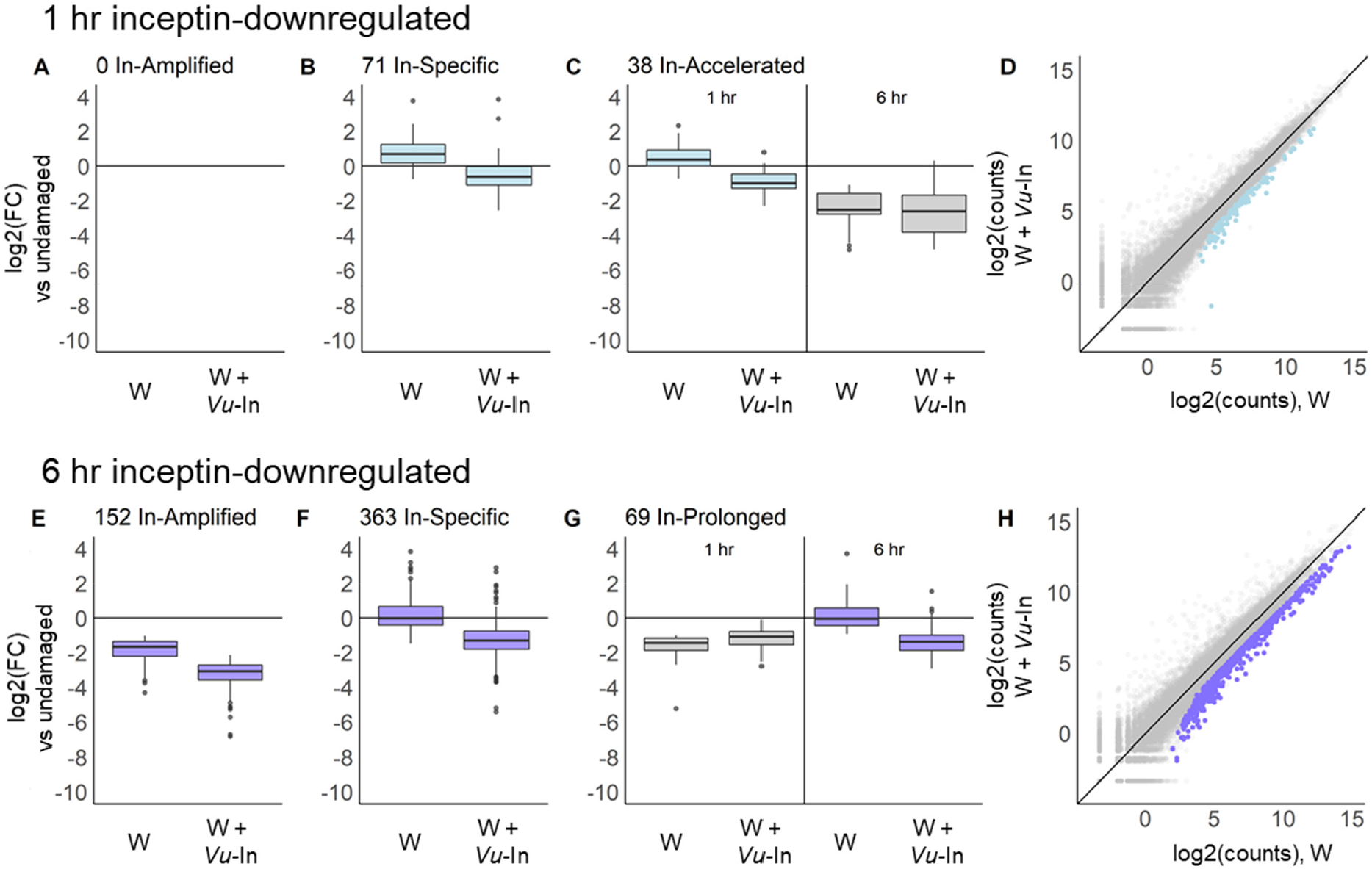
Transcriptional regulation of *Vu-*In-downregulated genes at 1 hr and 6 hr relative to wounding alone. Boxplots indicate magnitude fold change (FC) relative to undamaged treatment for categories of genes in **A**, *Vu*-In-Amplified; **B**, *Vu*-In-Specific; and **C**, *Vu*-In-Accelerated sets at 1 hr. **D**, Plot of log2(counts) after wounding (W) or W+ *Vu-*In for genes in **A**-**C**. Note that no genes fall in the category “*Vu-*In Amplified”. FC magnitude of **E**, *Vu*-In-Amplified; **F**, *Vu*-In-Specific; and **C**, *Vu*-In-Prolonged gene sets at 6 hr. **H**, Plot of log2(counts) after wounding (W) or W+ *Vu-*In for differentially expressed genes in **E**-**G**. Summary of filtering criteria for the *Vu-In* induced category assignments for specific, amplified, accelerated and prolonged groups are detailed in Fig. S3. For panels C and G, the behavior of same set of genes is shown at both 1 hr and 6 hr. Cowpea genes in each category underlying illustrated patterns are detailed in Tables S2 & S3.

**Fig. S5.**
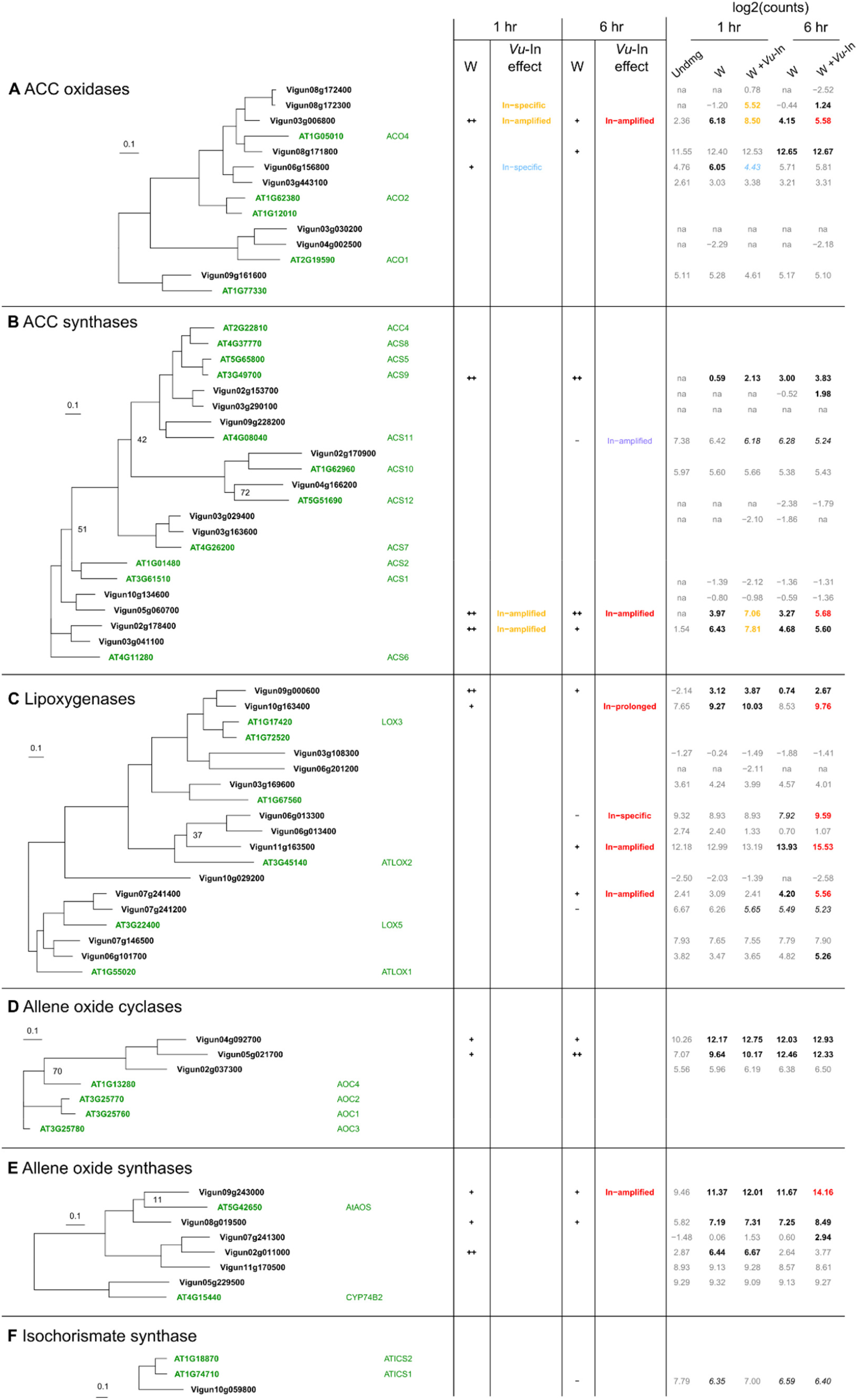
Wound- and *Vu-*In-induced changes in gene expression for defense hormone biosynthetic gene families. Phylogenetic trees display relationships between Arabidopsis and cowpea homologs of A, ACC oxidases, B, ACC synthases, C, lipoxygenases, D, Allene oxide cyclases (AOC), E, Allene oxide synthases (AOS), and F, Isochorismate synthases. Branch lengths indicate substitutions per site. Nodes with bootstrap labels indicate support in <70% of replicates. For all panels, cowpea homologs of Arabidopsis enzymes with known function were identified by TBLASTN search of the cowpea genome. Under column W, the effect of wounding is indicated for both timepoints relative to unwounded tissue at 1 hr. “+” indicates log2(FC) > 1 and Benjamini-Hochberg adjusted p<0.05, “++” indicates log2(FC) > 3 relative to unwounded tissue. Category of *Vu-*In-induced change is indicated for both timepoints (filters as in Fig. S3). Treatment average of log2-normalized counts is shown. Significant differences relative to undamaged tissue at 1 hr are displayed in bold (significantly upregulated) or italics (significantly downregulated), with colors indicating additional effect of *Vu-*In.

**Fig. S6.**
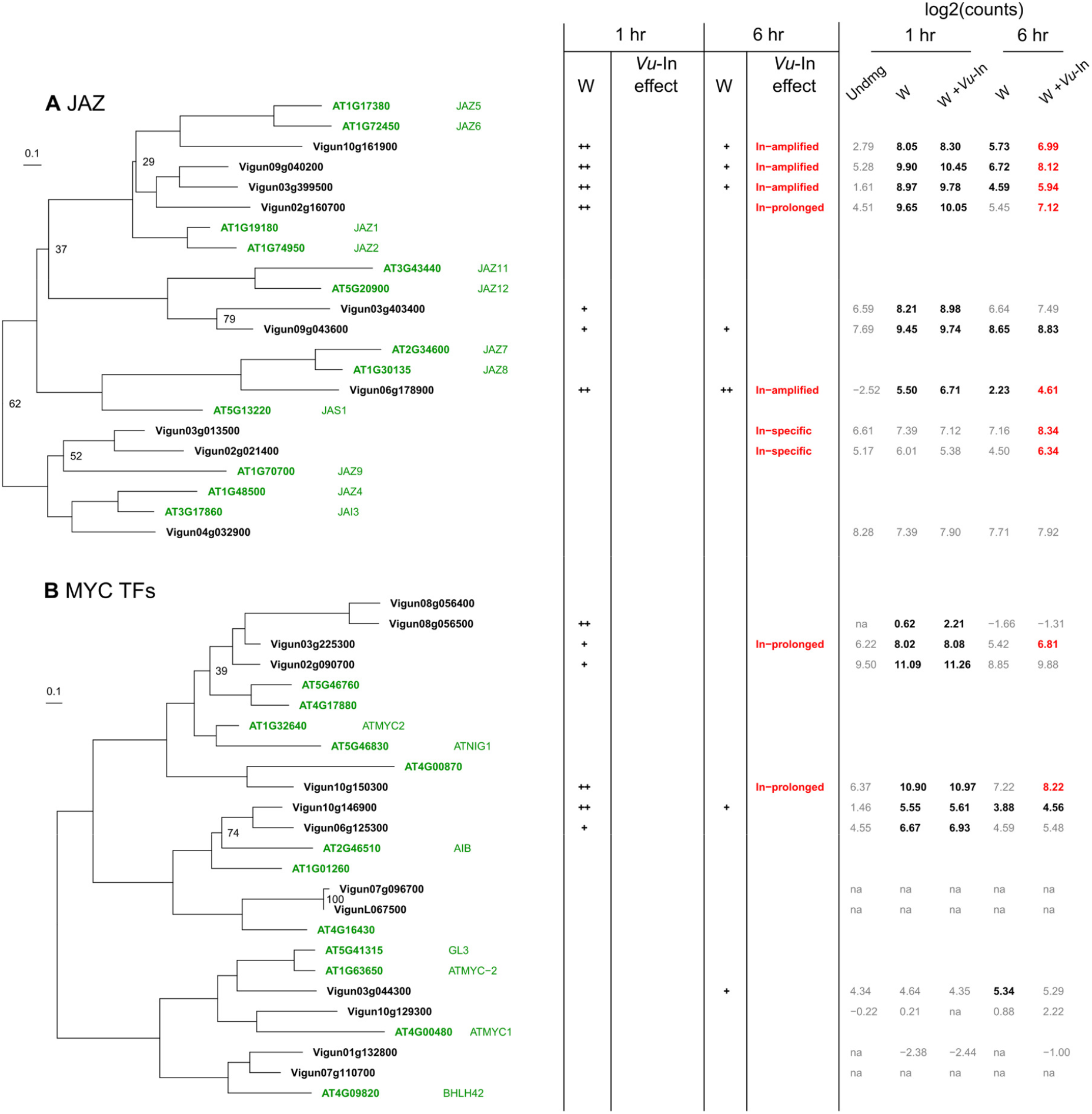
Wound- and *Vu-*In-induced changes in gene expression for gene families involved in jasmonate-related transcriptional regulation. Phylogenetic trees display relationships between Arabidopsis and cowpea homologs of A, JAZ repressors, B, MYC Transcription factors. Branch lengths indicate substitutions per site. All nodes shown have >80% bootstrap support unless noted. For all panels, cowpea homologs of Arabidopsis enzymes with known function were identified by TBLASTN search of the cowpea genome. Under column W, the effect of wounding is indicated for both timepoints relative to unwounded tissue at 1 hr. “+” indicates log2(FC) > 1 and Benjamini-Hochberg adjusted p<0.05, “++” indicates log2(FC) > 3 relative to unwounded tissue. Category of *Vu-*In-induced change is indicated for both timepoints (filters as in Fig. S3). Treatment average of log2-normalized counts is shown. Significant differences relative to undamaged tissue at 1 hr are displayed in bold (significantly upregulated) or italics (significantly downregulated), with colors indicating additional effect of *Vu-*In.

**Fig. S7.**
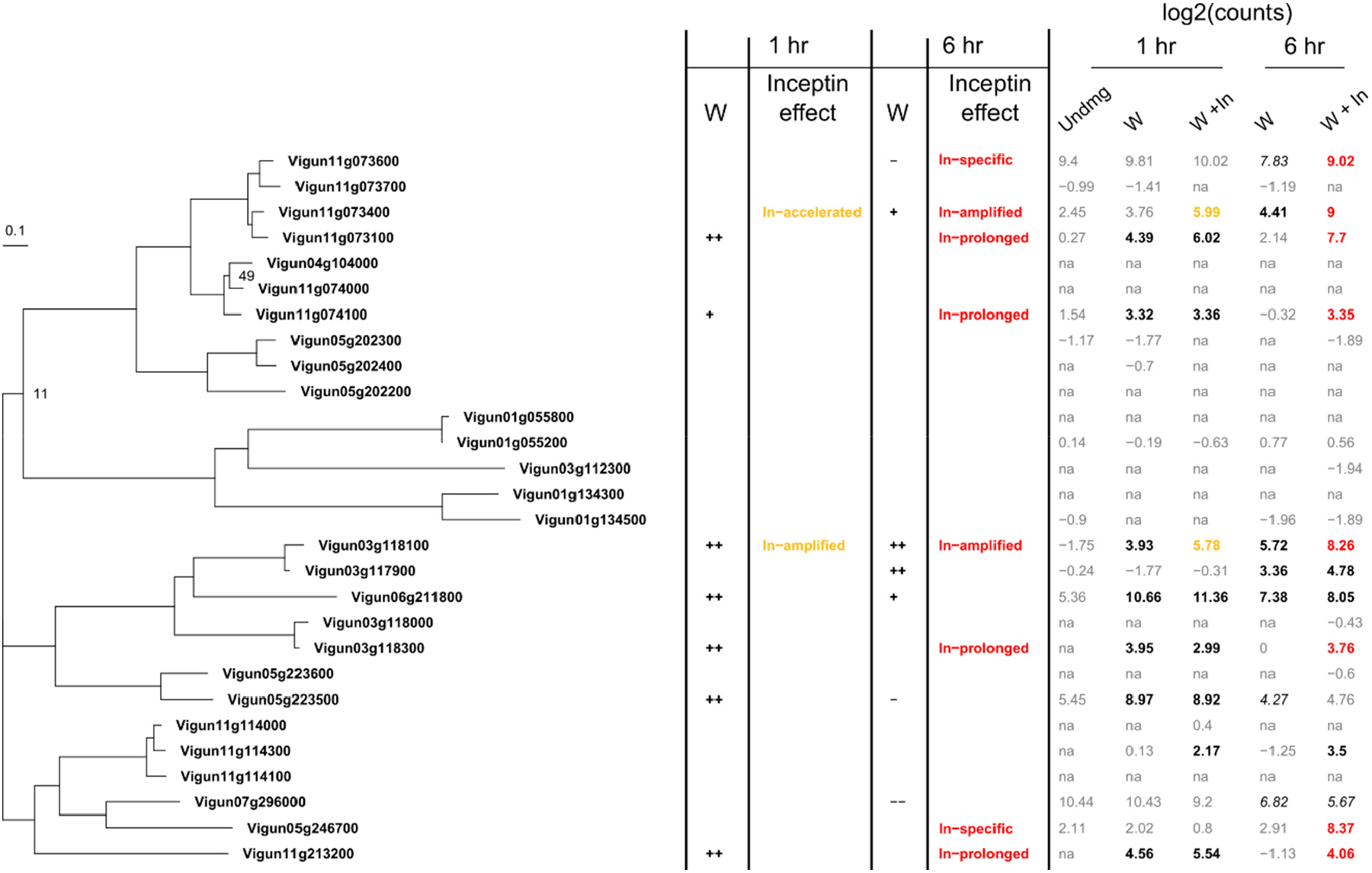
Wound- and *Vu-*In-induced changes in gene expression for terpene synthases (TPS). Branch lengths indicate substitutions per site. All nodes shown have >80% bootstrap support unless noted. Cowpea homologs were identified by TBLASTN using a Vigun05g246700 query sequence. Under column W, the effect of wounding is indicated for both timepoints relative to unwounded tissue at 1 hr. “+” indicates log2(FC) > 1 and Benjamini-Hochberg adjusted p<0.05, “++” indicates log2(FC) > 3 relative to unwounded tissue. Category of *Vu-*In-induced change is indicated for both timepoints (filters as in Fig. S3). Treatment average of log2-normalized counts is shown. Significant differences relative to undamaged tissue at 1 hr are displayed in bold (significantly upregulated) or italics (significantly downregulated), with colors indicating additional effect of *Vu-*In.

**Fig. S8.**
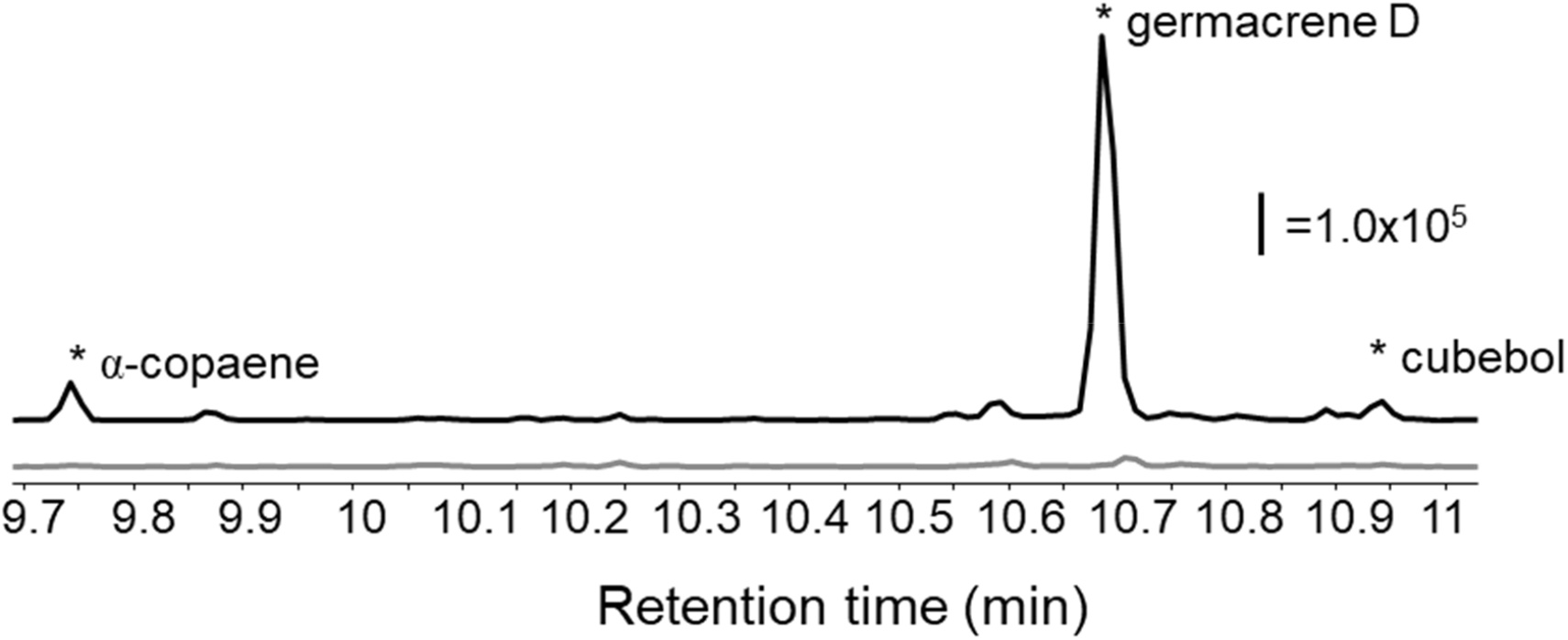
GC/MS total ion chromatogram (TIC) trace *N. benthamiana* volatile pools from leaf tissue extracts. Leaves were inoculated with *Agrobacterium* strains expressing empty vector or cowpea *VuTPS, Vigun11g073100*. Scale bar indicates arbitrary units of relative compound abundance for two treatments, TPS expression (black, top line) or empty vector (gray, bottom line).

**Fig. S9.**
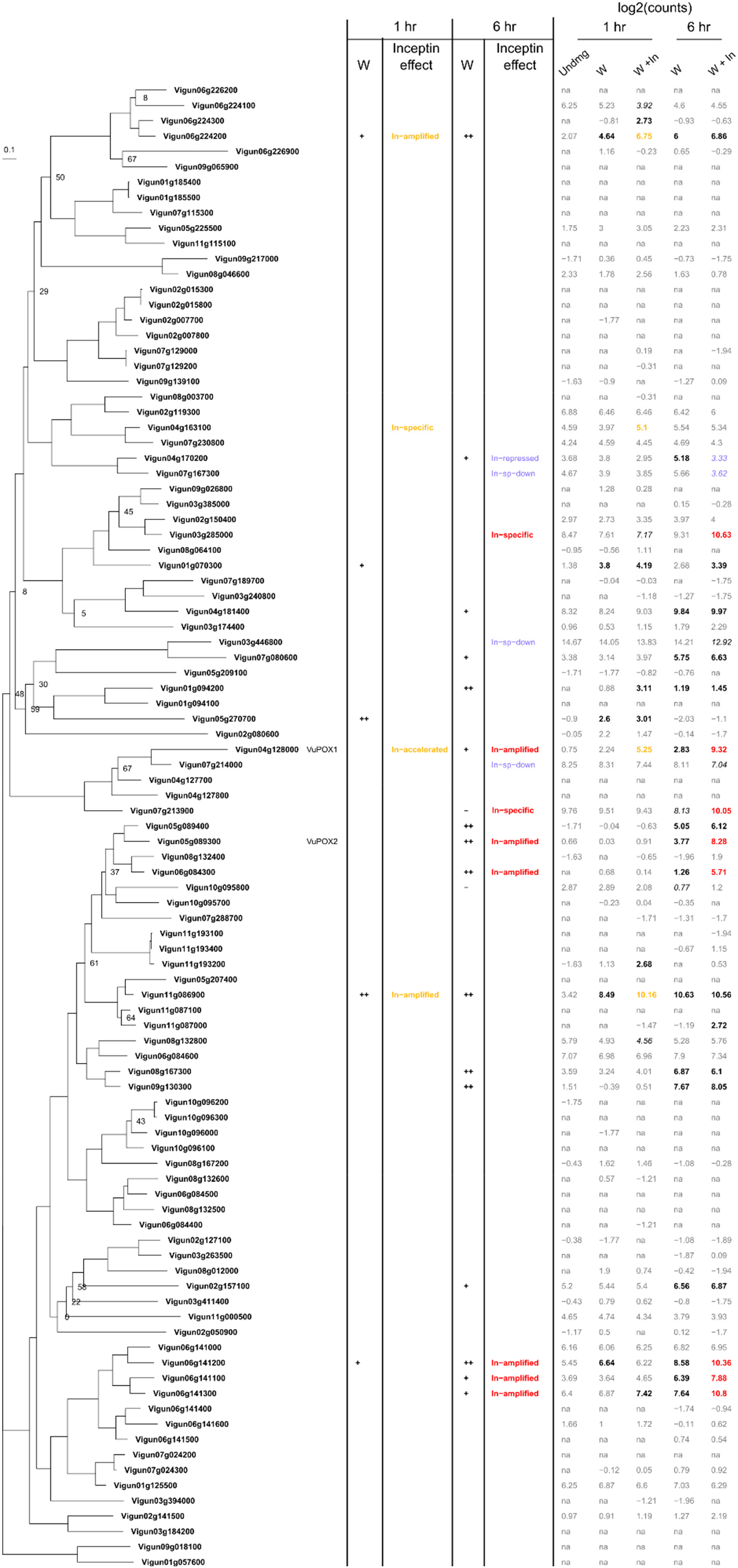
Wound- and *Vu-*In-induced changes in gene expression for peroxidases (POX). Branch lengths indicate substitutions per site. All nodes shown have >80% bootstrap support unless noted. Cowpea homologs were identified by TBLASTN search of the cowpea genome using a Vigun05g089300 query sequence. Under column W, the effect of wounding is indicated for both timepoints relative to unwounded tissue at 1 hr. “+” indicates log2(FC) > 1 and Benjamini-Hochberg adjusted p<0.05, “++” indicates log2(FC) > 3 relative to unwounded tissue. Category of *Vu-*In-induced change is indicated for both timepoints (filters as in Fig. S3). Treatment average of log2-normalized counts is shown. Significant differences relative to undamaged tissue at 1 hr are displayed in bold (significantly upregulated) or italics (significantly downregulated), with colors indicating additional effect of *Vu-*In.

**Fig. S10.**
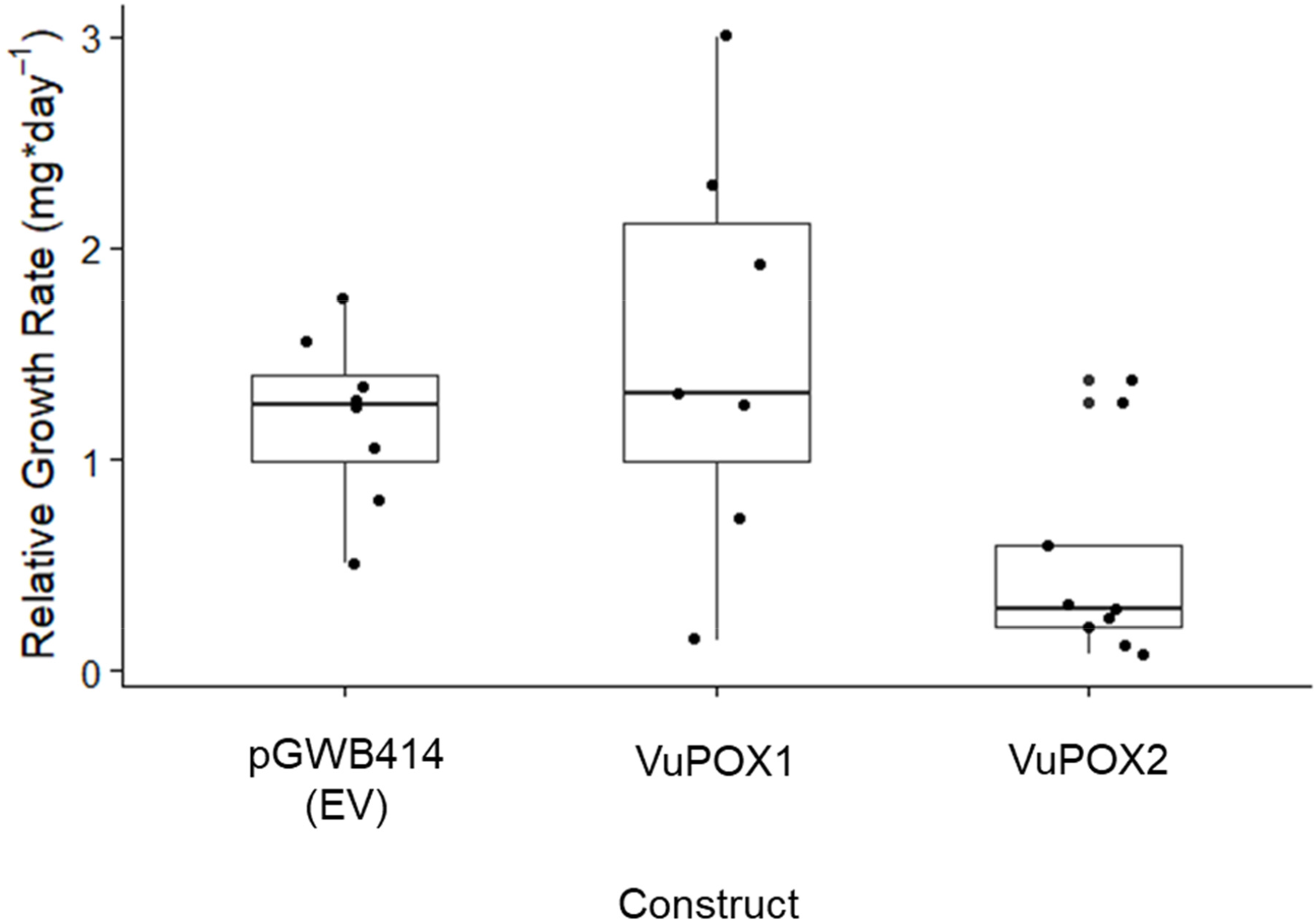
Expression of VuPOX2 but not VuPOX1 in *N. benthamiana* lowered *S. exigua* growth rates. 2nd instar larvae were caged with *N. benthamiana* leaf discs previously infiltrated with *Agrobacterium* strains to express either *VuPOX* gene or empty vector (EV). After 48 hr, caterpillars were weighed and relative growth rate was calculated.

